# Commensal Engineered Living Materials for Sustained Secretome Delivery and Wound Healing

**DOI:** 10.64898/2026.11.04.764344

**Authors:** Claire O. Lois, Elizabeth A. Grice, April M. Kloxin, Millicent O. Sullivan

**Affiliations:** Department of Chemical and Biomolecular Engineering, University of Delaware, _Newark_, DE, USA, _19716_; Department of Dermatology, University of Pennsylvania, Philadelphia, PA, USA, _19104_; Department of Materials Science and Engineering, University of Delaware, _Newark_, DE, USA, _19716_; Delaware Biotechnology Institute, _Newark_, DE, USA, _19713_; Department of Biomedical Engineering, University of Delaware, _Newark_, DE, USA, _19716_

## Abstract

Using living cells as dynamic biological factories offers immense potential for the continuous, localized production of therapeutic proteins within compromised tissues. This strategy is particularly compelling for treating chronic non-healing wounds, which lack therapeutics that are able to function effectively in the inflammatory microenvironment. Commensal bacteria have been shown to promote healing in these hostile microenvironments, yet applying live microbes directly onto compromised tissue is clinically unfeasible due off-target safety risks and engraftment challenges. Herein, we establish a well-defined commensal Engineered Living Material (cELM) that allows encapsulation, retention, function, and delivery of native beneficial bacteria for localized production and release of a pro-healing secretome over weeks. We specifically encapsulated a pro-healing human wound bacterial isolate, *Alcaligenes faecalis,* using biocompatible, click-reactive hydrogel building blocks (thiol- or vinyl-sulfone-functionalized multi-arm poly(ethylene glycol)) and systematically varied the density of the polymer network and of the encapsulated bacteria to establish design rules for creating pro-healing cELMs. Physical confinement in the hydrogel network ensured bacterial biocontainment, while the hydrogel mesh size still permitted diffusion of the therapeutic secretome. Notably, controlling encapsulation density revealed an inverse relationship wherein lower cell densities yielded significantly enhanced per-cell metabolic activity and provided sustained production of pro-healing secreted factors. In a murine diabetic wound model, these *A. faecalis* cELMs shielded the encapsulated bacteria from host clearance, maintained biocontainment, accelerated re-epithelialization, and improved wound bed maturity. Together, these findings establish design principles for cELMs that couple bacterial biocontainment with sustained secretome delivery, providing a promising platform for localized treatment of chronic wounds.

**One Sentence Summary:** Engineered living materials entrap therapeutic bacteria while sustaining their function and enabling secretome delivery for improved wound healing.

## INTRODUCTION

Engineered living materials (ELMs), in which viable cells are embedded within designed matrices, offer a promising strategy across diverse biomedical and biotechnological applications, from environmental sensing and biomanufacturing to tissue regeneration and targeted drug delivery (*1–8*). Integrating non-engineered commensal bacteria into ELMs offers a distinct opportunity to harness naturally evolved microbial functions, including the production of complex secretomes that support host tissue homeostasis and repair. However, current ELM approaches largely focus on a narrow range of model organisms, which are often genetically engineered for survival in engineered microenvironments (*1, 4, 9, 10*). Extending these approaches to non-engineered commensal species presents significant opportunities yet also introduces new challenges: *i)* maintaining non-engineered bacteria viability and function outside their native niches while ensuring physical biocontainment; and *ii)* sustaining production and release of bioactive secreted factors for therapeutic applications. Addressing these challenges requires understanding how the ELM microenvironment influences commensal bacterial survival, metabolic activity, and secretory function, as well as release of the resulting secretome. Establishing these relationships is essential for designing living materials that harness the native therapeutic potential of commensal bacteria.

One compelling application is the treatment of chronic non-healing wounds such as diabetic foot ulcers (DFUs), in which healing is stalled by hyper-inflammation and the overexpression of matrix metalloproteinases (MMPs) that degrade the extracellular matrix (ECM) and impair re-epithelialization (*11–22*). Bacteria in these wound microenvironments are often viewed indiscriminately as pathogens, with the clinical standard of care relying on broad-spectrum antimicrobials that eliminate both pathogenic and beneficial bacteria and unintentionally strip the wound bed of vital commensals that actively support tissue repair (*23–30*). Emerging evidence highlights the contributions of beneficial commensal species to tissue repair, including *Alcaligenes faecalis* (*A. faecalis*), a low-abundance human wound commensal (*31*). Direct topical application of live *A. faecalis* has been shown to promote healing in diabetic wounds through secretion of factors that downregulate MMP expression in keratinocytes, mitigating tissue degradation and restoring re-epithelialization (*11, 13, 15, 31, 32*). However, direct application of live bacteria into wounds raises concerns surrounding safety, reproducibility, and dose control (*2, 25, 26, 28*). Using encapsulation techniques to physically confine therapeutic microbes within a biocompatible matrix offers a potential strategy to overcome these limitations for safer, reproducible, and controllable microbe-based treatments, highlighting the need for ELMs that integrate and support non-engineered commensal bacteria.

To enable safe, reproducible ELMs encapsulating commensal species, the synthetic matrix must shield therapeutic bacteria from host defenses and rapid clearance, support sustained cell viability and secretory function, and permit the localized transport of bioactive secretome components (*1–4, 33–35*). Hydrogels provide structured, hydrated microenvironments in which network properties can be tuned to balance bacterial protection and confinement with biological function and molecular transport. In particular, poly(ethylene glycol) (PEG)-based hydrogels are well-defined, biocompatible materials widely used for the encapsulation of mammalian cells for both fundamental and applied studies (*36–38*), though with limited exploration of bacterial encapsulation (*39*). The tunable network properties in PEG-based hydrogels provide a means to investigate how physical confinement and mass transport influence commensal bacterial function and secretome release, including in the harsh microenvironment of the DFU (*34, 36, 37, 40, 41*). Here, we selected four-arm PEG functionalized with vinyl sulfone (PEG-VS4) or thiol (PEG-SH4) groups, which undergo thiol–Michael addition under mild aqueous conditions to form a covalently crosslinked network (*36, 40*). We hypothesized that these networks could physically confine *A. faecalis* while sustaining bacterial viability and function and permitting release of pro-healing secretome components. We further sought to determine how polymer network density and initial bacterial encapsulation density collectively influenced bacterial viability, metabolic activity, and secretome production and release.

In this work, we established a commensal engineered living material (cELM) platform designed for the sustained, localized delivery of the native *A. faecalis* secretome to promote diabetic wound healing. We systematically varied polymer network density (5–20 wt%) and bacterial encapsulation density (1 × 10⁸, 5 × 10⁸, and 9 × 10⁸ CFU/mL) and characterized the retention, spatial organization, and long-term metabolic viability of *A. faecalis* within bulk hydrogels. We identified an optimal 10 wt% hydrogel construct that maintained >90% cell viability and complete physical biocontainment over 14 days, a timeframe over which *A. faecalis* in suspension previously had been shown to promote wound healing (*42*). The kinetic release profiles and composition of the soluble protein secretome components were validated quantitatively. In addition, the insoluble secreted matrix was assessed qualitatively with staining and imaging, suggesting the formation of biofilm-like structures. Furthermore, the functional biological activity of the released secretome was evaluated *in vitro* using a primary human epithelial keratinocyte scratch closure assay. Counterintuitively, lower bacterial encapsulation densities 1 × 10⁸ CFU/mL) yielded an enhancement in per-cell metabolic output and superior secretome release, which significantly promoted human keratinocyte migration *in vitro*. The viability and function of the *A. faecalis* cELM then was evaluated *in vivo* using a preclinical murine diabetic wound model, where cELMs prevented host clearance of encapsulated cells, maintained absolute biocontainment in the wound bed, and extended significant wound closure acceleration through the full 14-day treatment period relative to unencapsulated suspension controls. Together, these studies establish relationships amongst material properties, bacterial loading and function, and secretome delivery, providing design principles for biocontained living materials that harness native commensal functions to promote tissue repair.

## RESULTS

### Influence of network density on encapsulated *A. faecalis* viability and metabolism

To evaluate the potential of commensal bacteria-laden hydrogels as biocontained, secretome-producing therapeutics, we first investigated how PEG hydrogel network density influenced the viability and metabolic activity of encapsulated *A. faecalis*. We fabricated bulk hydrogels at a fixed initial cell density of 5 × 10⁸ CFU/mL, guided by bacterial suspension concentrations previously shown to promote diabetic wound healing (*31*), while varying total weight percentages of PEG (5, 10, and 20 wt%) to modulate network density. These formulations yielded estimated mesh sizes of approximately 7.2 nm, 4.8 nm, and 2.7 nm, respectively, based on the Flory-Rehner Equation as described in the Methods (**Figure 1A**). We selected 5 kDa PEG-VS4 and PEG-SH4 as the building blocks based on our prior work demonstrating the utility of these constructs for forming well-defined, tunable hydrogel networks suitable for biomolecule encapsulation and release (*41*).

**Figure 1.**
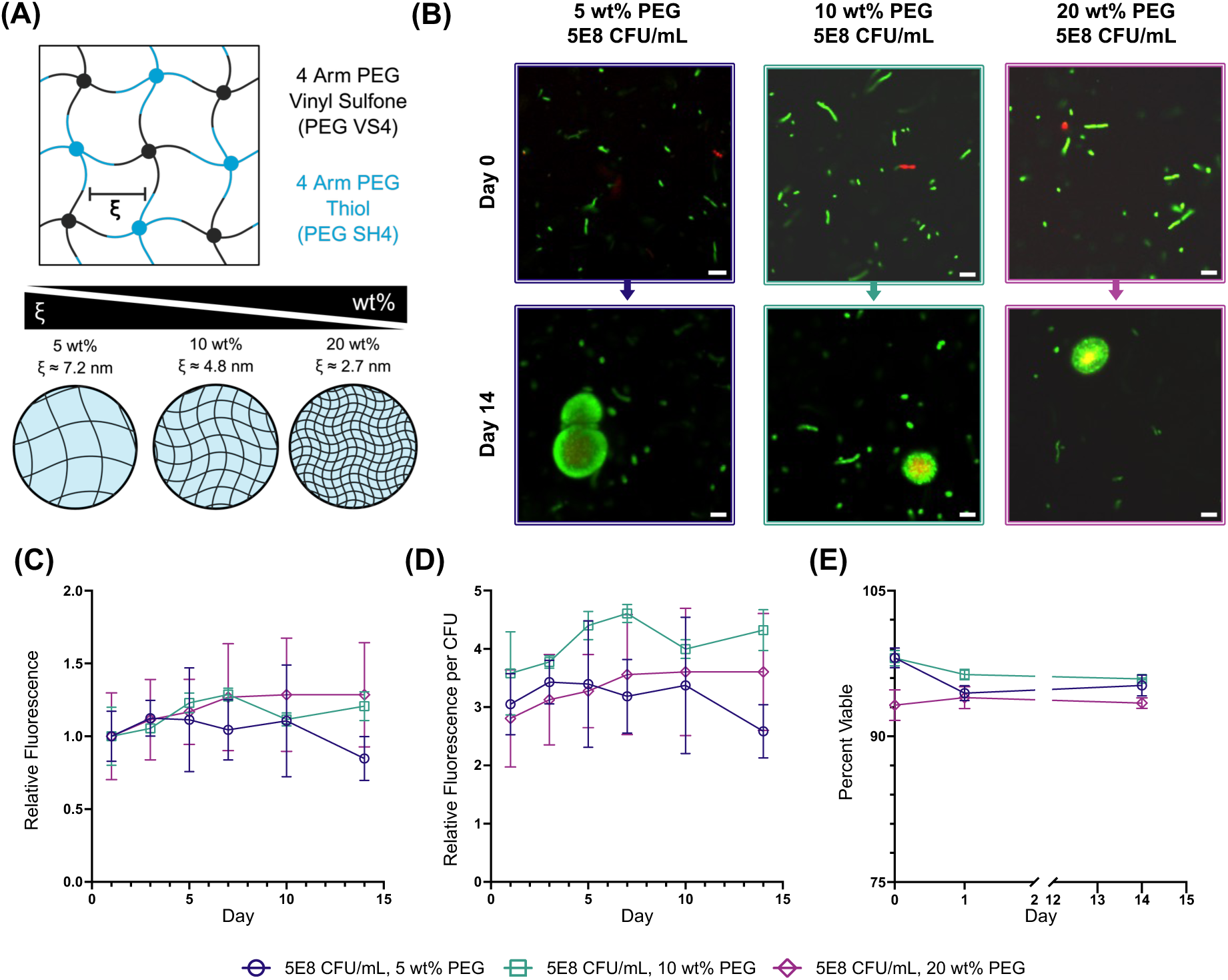
Characterization of polymer weight percent variation on network mesh parameters and encapsulated bacterial viability. (A) Schematic detailing the theoretical inverse scaling relationship between polymer weight percentage (5 wt%, 10 wt%, and 20 wt%) and estimated network mesh size derived via the Flory-Rehner equation. (B) Live/Dead imaging of cells encapsulated in the various hydrogels (viable cells = green, membrane-compromised cells = red) immediately following encapsulation (day 0) and after 14 days of *in vitro* incubation. Scale bar = 5 µm. (C) Cumulative metabolic activity profiles of whole, intact hydrogels tracked over 14 days via PrestoBlue fluorescence assays, normalized to day 1. (D) Relative metabolic fluorescence outputs normalized directly per individual encapsulated colony-forming unit (CFU). (E) Longitudinal quantification of percentage cell viability determined via Imaris. Data represent mean (N=4) ± standard deviation (SD).

We used confocal microscopy to track the three-dimensional (3D) spatial distribution and viability of bacteria cells encapsulated within the different hydrogel formulations (5, 10, and 20 wt%), staining cells in the 3D constructs with SYTO 9 (Live) and propidium iodide (PI, Dead) over 14 days (**Figure 1B**). Qualitatively, encapsulated cells showed a uniform distribution on day 0, with staining patterns that corresponded to the expected dimensions of encapsulated single cells. By day 14, some encapsulated bacterial cells appeared larger, with halo-like staining patterns of ∼5 – 10 µm in diameter around individual cells. Although bacterial counts remained relatively constant (**Figure 1E)**, these enlarged fluorescent regions may reflect the accumulation of extracellular nucleic acids or other bacterial-associated material, potentially consistent with the formation of a biofilm-like matrix. Biofilm-like matrices often contain nucleic acid fragments, which are known to become stained by SYTO 9 and PI (*43, 44*).

Quantitatively, we assessed total metabolic activity using the non-destructive PrestoBlue metabolic activity reagent to analyze the supernatant over the 14-day tracking window. The average total fluorescence from each gel was similar as a function of time when compared across the three PEG densities (**Figure 1C**). Assessment of hydrogel swelling over 14 days revealed maintained matrix properties. When normalized to unladen blank hydrogels, volumetric swelling ratios confirmed uniform structural stability across all cell polymer densities (**Figure S1**). When we normalized the metabolic output per colony-forming unit (CFU), all three formulations also exhibited similar and relatively constant profiles over time, suggesting the maintenance of cell viability given the constant bacterial cell counts. The 10 wt% network exhibited a slight elevation in normalized metabolic efficiency by day 7 (**Figure 1D**). Analysis of the live / dead confocal images confirmed that the percentage of viable cells remained consistently above 90% across all weight percentages throughout the 14 days (**Figure 1E**).

We selected the 10 wt% formulation for subsequent studies as an intermediate composition intended to balance hydrogel network integrity with molecular transport. Indeed, previous studies of 4-arm PEG hydrogels have demonstrated that polymer concentration influences network connectivity and structural homogeneity, with dilute precursor conditions potentially promoting incomplete crosslinking, dangling chains, and other network defects (*45, 46*). Consistent with these architectural limits, 5 wt% hydrogels exhibited detectable bacterial cell escape over extended culture (**Figure S2**). Conversely, increasing polymer concentration reduces hydrogel mesh size and can restrict macromolecular diffusion through the network (*47*), as demonstrated in our prior work (*36*). Given the modestly higher metabolic activity observed for bacteria encapsulated in the 10 wt% formulation and its intermediate estimated mesh size that should be permissive to protein diffusion (**Table S1**), we selected 10 wt% composition to support bacterial entrapment within a crosslinked hydrogel network while facilitating the exchange of nutrients, metabolites, and bacterially secreted therapeutic factors.

### Impact of initial encapsulation density on long-term bacterial behavior and viability

We next sought to test whether bacterial encapsulation density would alter bacterial viability, secretome output, and intra-gel matrix phenotype, hypothesizing that changes to spatial confinement and cell proximity would affect cell behavior. As expected, increasing the initial seeding density (1 × 10⁸, 5 × 10⁸, and 9 × 10⁸ CFU/mL) systematically decreased the average inter-bacterial distance (*d*) within the 10 wt% hydrogel network (**Figure 2A**). Longitudinal monitoring of hydrogel swelling demonstrated stable network properties over 14 days. Volumetric swelling ratios normalized to corresponding unladen blank hydrogels showed consistent structural stability across varying cell encapsulation densities (**Figure S1**). Confocal imaging of the resulting SYTO 9 and PI-stained cELMs revealed distinct halo-like formations, consistent with our observations with 5 × 10⁸ CFU/mL in different wt% gels (**Figure 1)**. Interestingly, at the lowest bacterial encapsulation density (1 × 10⁸ CFU/mL), halo-like structures appeared larger and more clearly defined than at higher bacterial encapsulation densities after 14 days, potentially reflecting differences in the local distribution and accumulation of nucleic acid-containing material, including extracellular DNA that may be associated with biofilm formation (**Figure 2B**).

**Figure 2.**
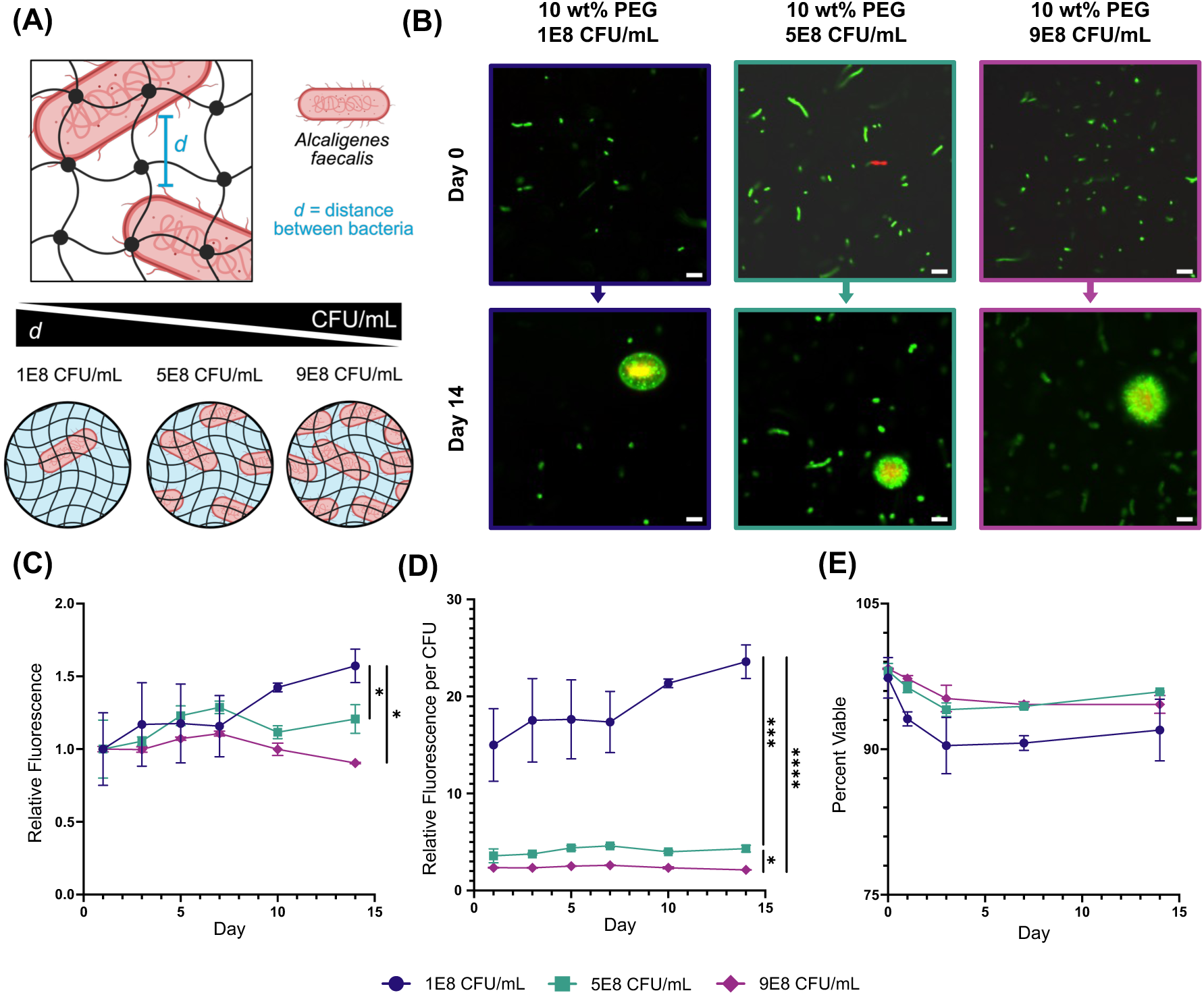
Influence of initial bacterial encapsulation density on viability and metabolic activity. (A) Schematic illustrating the relationship between encapsulation density and the resultant average spatial distance (d) separating encapsulated *A. faecalis* cells within a baseline 10 wt% PEG hydrogel network. (B) Representative Live/Dead imaging of cells encapsulated within hydrogels at the time of encapsulation and after 14 days. Scale bar = 5 µm. (C) Cumulative metabolic activity profiles of whole, intact hydrogels tracked over 14 days via PrestoBlue fluorescence assays, normalized to day 1. (D Relative metabolic fluorescence outputs normalized directly per individual encapsulated colony-forming unit (CFU). (E) Longitudinal quantification of percentage cell viability determined via Imaris. Data represent mean (N=4) ± standard deviation (SD). For all plots: *P ≤ 0.05; **P ≤ 0.01; ***P ≤ 0.001, ****P ≤ 0.0001 by 2-Way ANOVA with Tukey’s multiple comparison test.

The metabolic profiles for each condition also reflected a phenotypic divergence as a function of cell density. Hydrogel samples laden with 1 × 10⁸ CFU/mL demonstrated a significant, steady increase in overall relative metabolic fluorescence over 14 days, whereas the metabolic fluorescence plateaued or slightly declined after day 7 in cELMs at higher cell densities (**Figure 2C**). This trend became highly pronounced when we normalized metabolic activity per CFU. While raw metabolic activity readouts per gel remained similar, normalization on a per-CFU basis revealed that cells in the 1 × 10⁸ CFU/mL condition were on average ∼ 5 times more active than cells in gels containing 5 × 10⁸ CFU/mL and on average ∼ 8 times more active than cells in gels containing 9 × 10⁸ CFU/mL (**Figure 2D**). This observation may indicate a significant metabolic advantage and minimized nutrient competition at lower initial bacterial encapsulation densities. Despite these clear phenotypic and metabolic differences, overall cell viability remained statistically uniform and high across all density groups, with all cohorts maintaining about 90% viability through day 14 (**Figure 2E**).

### Evaluation of intra-matrix biofilm-like secretions over time

Prompted by the distinct staining patterns observed during confocal imaging noted above, we examined the intra-gel accumulation and temporal development of biofilm-like secretome components produced by encapsulated *A. faecalis* (**Figure 3A**) (*43, 44, 48*). We harvested the hydrogel samples at designated timepoints and then fixed and stained the encapsulated cells with a 0.1% crystal violet (CV) solution to label the accumulated protein-polysaccharide matrix. Quantitative colorimetric analysis of the resolubilized CV dye revealed a steady, time-dependent increase in absorbance at 590 nm across all cell-laden groups over the 14-day incubation period, confirming active deposition of proteins and biofilm-like matrix (**Figure 3B**). Paralleling the metabolic activity trend, the 1 × 10⁸ CFU/mL cohort exhibited the sharpest increase in accumulation, achieving a significantly higher total absorbance by day 14 compared to higher encapsulation densities.

**Figure 3.**
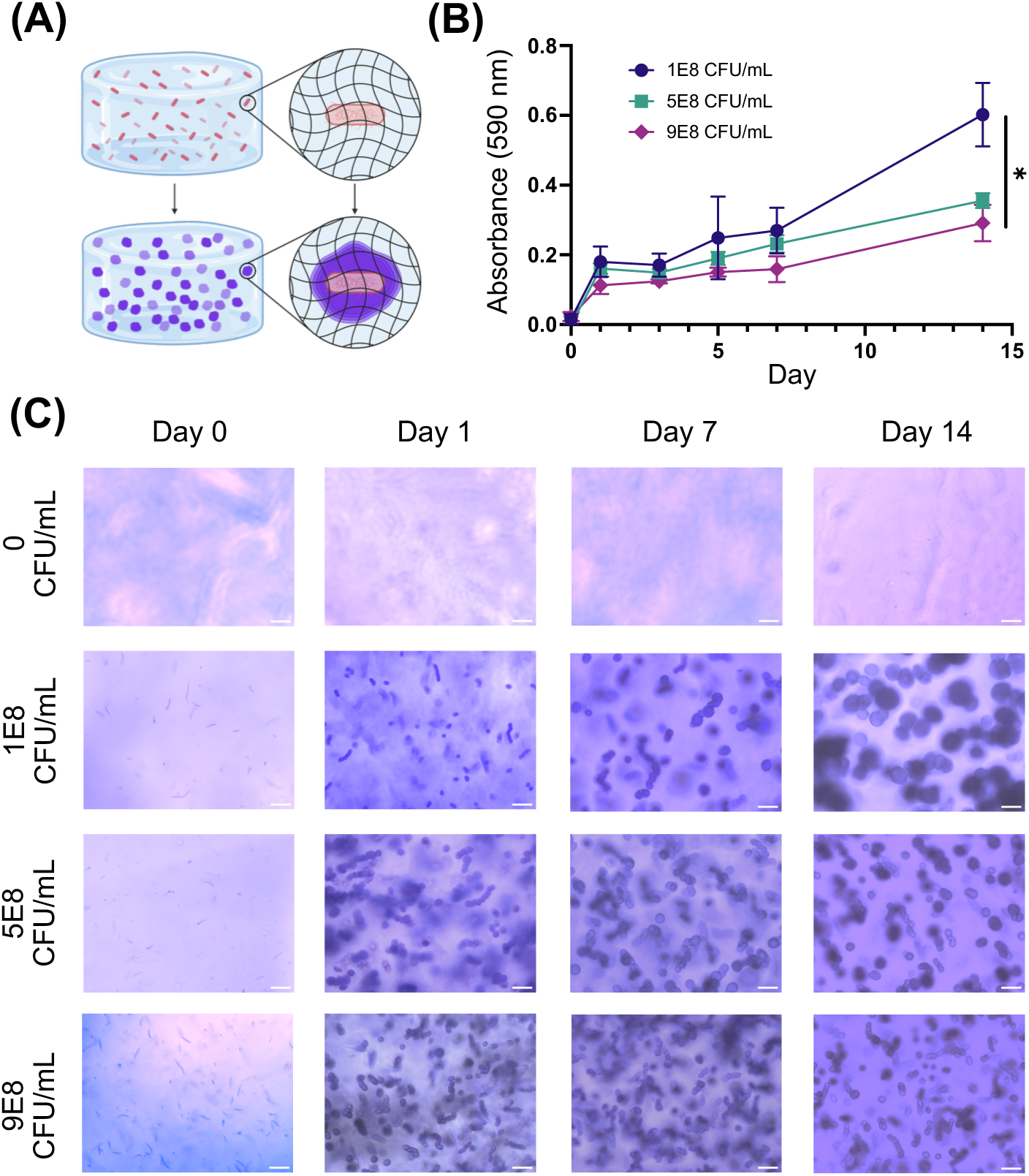
Evaluation of intra-gel biofilm-like matrix deposition. (A) Schematic of stained hydrogels showing the hypothesized phenomena. (B) Quantitative colorimetric tracking of intra-gel matrix accumulation over 14 days, evaluated by absorbance measurements of resolubilized crystal violet dye in 70% ethanol. Data represent mean (N=4) ± standard deviation (SD). *P ≤ 0.05 by 2-Way ANOVA with Tukey’s multiple comparison test. (C) Representative brightfield microscopy images intact hydrogels fixed and stained with 0.1% crystal violet across varying encapsulation densities on days 0, 1, 7, and 14, highlighting progressive formation/accumulation of biofilm-like matrix.

We also evaluated these CV-stained samples qualitatively via brightfield microscopy prior to dye solubilization (**Figure 3C**). Control hydrogels without encapsulated bacteria (0 CFU/mL) showed minimal non-specific dye retention over 14 days, retaining the same slight uniform color and full transparency over time. In contrast, cell-laden hydrogels demonstrated progressive, deep-purple aggregate formation, consistent with the morphologically similar features observed via confocal microscopy of SYTO9/PI-stained samples in **Figures 1** and **2**. While images from CV-stained samples (**Figure 3**) visually appeared to exhibit a higher density of aggregate formation than the SYTO9/PI-stained images for corresponding samples (**Figures 1** and **2**), the SYTO9-PI images represent a single slice in the z stack, whereas CV-stained images in **Figure 3** were taken through the entire volume of the hydrogel using a brightfield optical microscope. When CV-stained images were compared across different cell densities, an inverse relationship was apparent between aggregate size and cell density, with the size of the aggregate features appearing to decrease as encapsulation density increased (**Figure 3**). Representative three-dimensional reconstructions of the hydrogel constructs at days 0 and 14 further illustrate the spatial distribution of encapsulated bacteria and provide a qualitative assessment of changes in fluorescence over the incubation period (**Figure S3**).

### Characterization and composition of the secreted protein kinetics and diffusion

To further probe the secretory properties of encapsulated bacteria and assess the potential of the *A. faecalis* cELM as a therapeutic delivery platform, we characterized secreted protein release profiles from the hydrogels. Total protein in the supernatant of each hydrogel sample was quantified using the Pierce Rapid Gold bicinchoninic acid (BCA) protein assay. Building on our initial evaluation of hydrogel compositions, we first compared across the 5, 10, and 20 wt% PEG formulations at a fixed bacterial encapsulation density (5 × 10⁸ CFU/mL). The 5 wt% PEG hydrogels exhibited the highest cumulative protein release over 14 days on a per-gel basis, whereas the 10 wt% and 20 wt% hydrogels exhibited lower, more controlled cumulative release profiles (**Figure 4A**). These differences were consistent with the progressively smaller estimated mesh sizes of these formulations and thereby the greater potential for hindered diffusion in the more densely crosslinked networks. Normalizing cumulative protein release on a per-CFU basis yielded a similar trend (**Figure 4B**).

**Figure 4.**
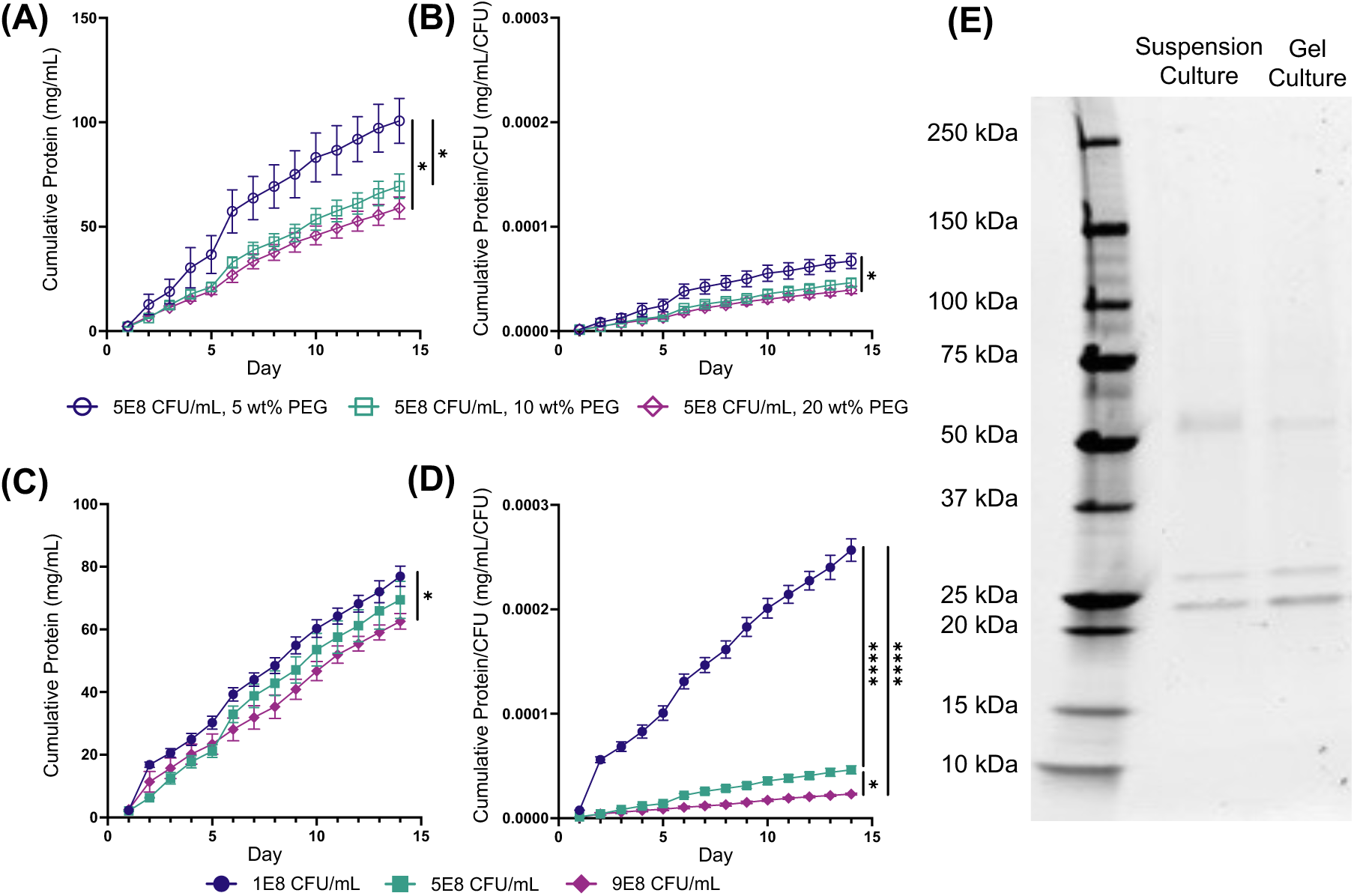
Protein secretion from hydrogels laden with bacteria. (A) Cumulative absolute protein accumulation kinetics and (B) corresponding per-CFU normalized protein secretion profiles released into the surrounding supernatant over 14 days from hydrogels containing a fixed bacterial density (5E8 CFU/mL) across varying polymer weight percentages (5, 10, and 20 wt%). (C) Cumulative absolute protein accumulation (D) and per-CFU normalized protein secretion curves released from 10 wt% PEG hydrogels across varying initial encapsulation densities (1E8, 5E8, and 9E8 CFU/mL). All protein concentrations were quantified via Pierce Rapid Gold BCA microplate assays under continuous sink conditions. (E) Representative sodium dodecyl sulfate-polyacrylamide gel electrophoresis (SDS-PAGE) separation profile stained with SYPRO Ruby, comparing the molecular weight distribution of the diffused protein secretome collected from 10 wt% gels containing 1E8 CFU/mL against raw conditioned media harvested from unencapsulated *A. faecalis* suspension cultures. For all plots: *P ≤ 0.05; **P ≤ 0.01; ***P ≤ 0.001, ****P ≤ 0.0001 by 2-Way ANOVA with Tukey’s multiple comparison test. Data represent mean (N=4) ± standard deviation (SD).

We next investigated the influence of bacterial encapsulation density (1 × 10⁸, 5 × 10⁸, 9 × 10⁸ CFU/mL) on protein release from the selected 10 wt% PEG formulation. Interestingly, hydrogels prepared at the lowest initial encapsulation density (1 × 10⁸ CFU/mL) exhibited the highest cumulative protein release into the supernatant, exceeding that of both the 5 × 10⁸ CFU/mL and 9 × 10⁸ CFU/mL conditions on a per-gel basis over 14 days (**Figure 4C**). Normalizing on a per-CFU basis further emphasized this difference, with the lowest cell density exhibiting significantly greater cumulative protein release per cell relative to higher-density formulations (**Figure 4D**). These findings are aligned with the differences observed in extracellular matrix accumulation (**Figure 3**) and bacterial metabolic activity (**Figure 2**), suggesting that bacterial encapsulation density influenced the accumulation and release of extracellular products. Additionally, we analyzed the molecular weight distribution of the released secreted proteins using sodium dodecyl sulfate-polyacrylamide gel electrophoresis (SDS-PAGE) stained with SYPRO Ruby (**Figure 4E**). The protein banding pattern of supernatants from the 10 wt% hydrogels with 1 × 10⁸ CFU/mL (pooled over 14 days) matched conditioned media harvested from unencapsulated

*A. faecalis* liquid suspension cultures after 18 hours of culture, with distinct bands at approximately 22–28 kDa and 55 kDa in both conditions (**Figure S4**). These data suggested that hydrogel encapsulation permitted the release of major soluble secreted proteins.

### *In vitro* functional potency of the encapsulated *A. faecalis* secretome on keratinocytes

We next evaluated the biological activity of the secretome released from the 10 wt% formulation laden with low or medium densities of *A. faecalis*, reasoning that these cELM formulation conditions offered the most interesting potential for wound healing given the higher per-CFU protein release described above. Released secretome bioactivity was evaluated using an *in vitro* scratch closure assay with confluent primary human epithelial keratinocyte monolayers (**Figure 5A**) (*49*). A challenge arose when designing this experiment to appropriately control for cell numbers when comparing between the constant population of encapsulated bacteria in hydrogel cultures and the expanding population of bacteria in control liquid suspension cultures. To address this, suspension timepoints were selected by measuring time-integrated cell numbers over 24 hours (area under the curve, AUC), using cell counts within a representative 50 µL volume (**Figure S5**). Integrating growing suspension cell counts over 0–9 h yielded time-integrated cell numbers (AUC = 1.03E8) comparable to the constant 1 × 10⁸ CFU/mL hydrogel condition over 24 h (AUC = 1.20E8), whereas 0–18 h (AUC = 6.63E8) was comparable to the constant 5 × 10⁸ CFU/mL condition over 24 h (AUC = 6.0E8). Consequently, the 9 h and 18 h suspension time points were selected as matched controls for subsequent analyses (**Figure S5**).

**Figure 5.**
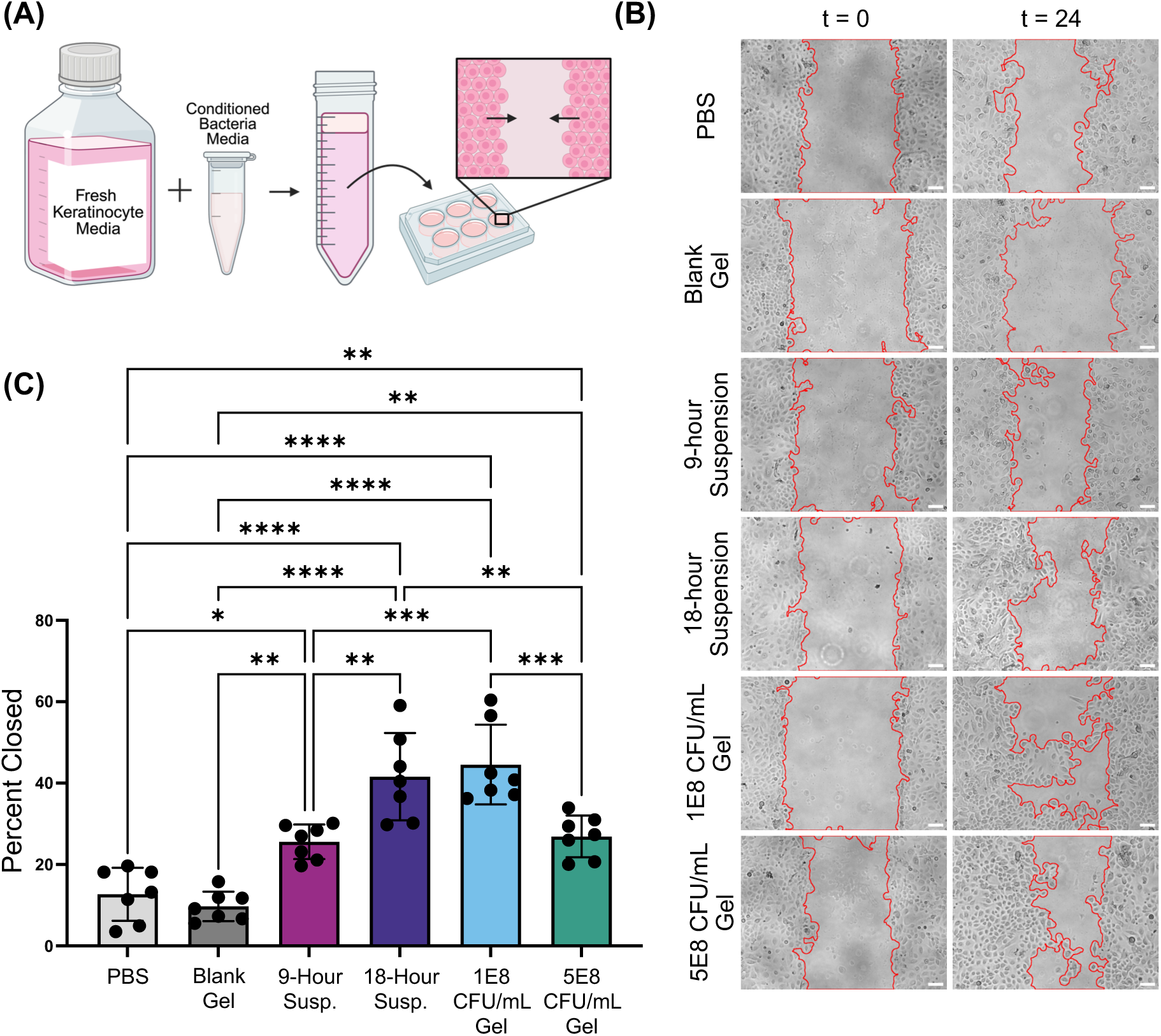
*In vitro* healing using keratinocyte scratch assays. (A) Conditioned media formation and treatment to keratinocyte scratches. (B) Images of keratinocyte scratches treated with 10% (v/v) of conditioned media as labeled. Images were taken at t=0 when scratches were made and t=24 corresponding to 24 hours of incubation with conditioned media. (C) Percent of original scratch closed after 24 hours incubation. For all plots: *P ≤ 0.05; **P ≤ 0.01; ***P ≤ 0.001, ****P ≤ 0.0001 by One-Way ANOVA with Tukey’s multiple comparison test. Data represent mean (N=7) ± standard deviation (SD).

In evaluating the bioactivity of the resulting supernatants, treatment of scratches with conditioned media from low-density *A. faecalis* cELMs (1 × 10⁸ CFU/mL) enhanced 24-hour scratch closure (36.6 ± 3.4%), with a statistically significant enhancement over both vehicle controls (cell-free PBS) (12.1 + 6.7%) and blank hydrogel (10.2 ± 5.2%), supporting bioactivity of the released secretome (**Figure 5C**). Similarly, conditioned media from mid-density *A. faecalis* cELMs (5 × 10⁸ CFU/mL) drove closure (33.5 ± 6.0%), significantly outperforming PBS and blank gel controls. Though not significant, comparison between the two hydrogels suggested a trend that the lower-density *A. faecalis* cELM promoted superior keratinocyte migration in comparison to the mid-density *A. faecalis* cELM, consistent with the inverse density-dependent secretome output. Conditioned media from the 18-hour suspension culture significantly enhanced scratch closure relative to media from the 9-hour suspension culture, consistent with greater accumulation of bioactive factors over the longer culture period.

Interestingly, despite being matched by time-integrated bacterial cell counts (Figure S2), the 1 × 10⁸ CFU/mL hydrogel and the 9-hour suspension culture conditions exhibited significantly different scratch closure, as did the 5 × 10⁸ CFU/mL hydrogel and the 18-hour suspension culture conditions (**Figure 5C**). Instead, the functional outcomes revealed the opposite pairing: scratch closure did not differ significantly between the 1 × 10⁸ CFU/mL hydrogel and the 18-hour suspension conditions, nor between the 5 × 10⁸ CFU/mL hydrogel and the 9-hour suspension conditions. These observations suggest that bacterial encapsulation density influences the released secretome in ways not predicted solely by cell numbers, underscoring the heightened per-cell metabolic and secretory output of the lower-density *A. faecalis* cELM. Importantly, these data demonstrate that hydrogel-encapsulated *A. faecalis* can produce and release a bioactive, pro-healing secretome that promotes human keratinocyte scratch closure.

### *In vivo* persistence, viability, and absolute biocontainment of live bacterial hydrogels

A central challenge in translating commensal bacteria-laden hydrogels into living therapeutic platforms is retaining viable bacteria at the treatment site for durations sufficient to support wound healing. Having established sustained bacterial viability, function, and bioactive secretome release *in vitro*, we next evaluated if an *A. faecalis* cELM could promote cell survival and wound healing *in vivo*. We chose an 8-mm full-thickness excisional wound model using 8-week-old diabetic (*db/db*) model mice, which exhibit impaired wound healing and provide a relevant model for investigating therapeutic strategies for human diabetic wounds. We first focused on evaluating bacterial persistence and physical biocontainment (**Figure 6A**). We treated wounds with an *A. faecalis*-laden cELM (5 × 10⁸ CFU/mL) or an equivalent dose of *A. faecalis* suspended in sterile saline solution, with the bacterial dose selected to align with prior studies investigating *A. faecalis* in diabetic wounds (*31*).

**Figure 6.**
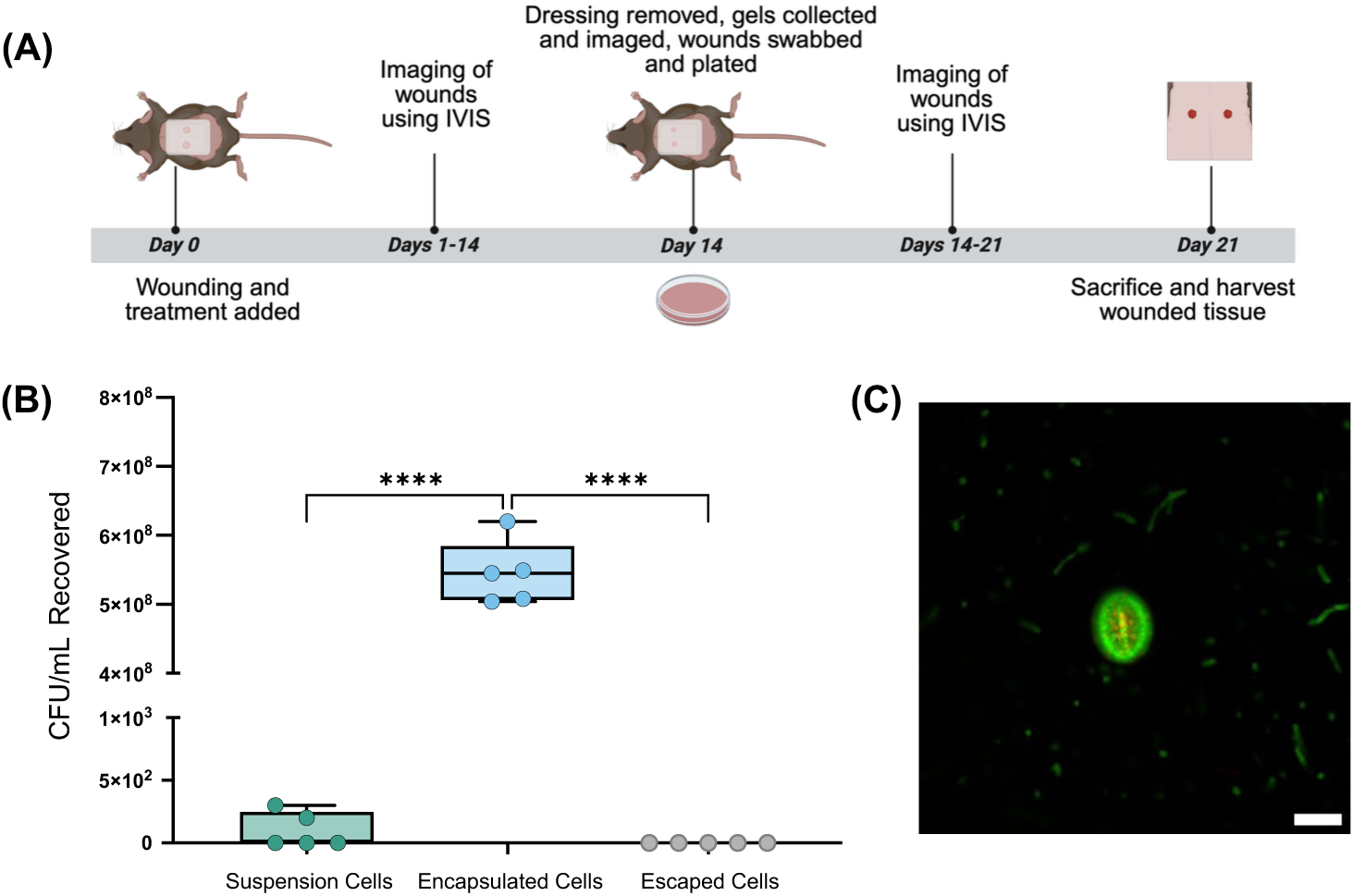
*In vivo* persistence, viability, and physical biocontainment of bacteria in living hydrogels in a diabetic murine model. (A) Experimental timeline outlining events of wounding, treatment removal, and evaluation of cell survival. (B) Localized bacterial recovery profiles quantified via colony-forming unit (CFU/mL) plating on nutrient agar at day 14. Groups compare topically applied free suspension cells, active cells encapsulated within the hydrogels, and surface wound swabs (escaped cells) taken from the surrounding tissue beds, establishing physical biocontainment. Data points represent individual values. (C) Representative image of hydrogel removed from wound and imaged using confocal microscopy. For all plots: ****P ≤ 0.0001 by One-Way ANOVA with Tukey’s multiple comparison test.

After 14 days, treatments were removed to assess bacterial persistence and local biocontainment. Notably, swabs collected from wounds treated with the free *A. faecalis* suspension yielded few to no recoverable bacterial colonies, whereas hydrogels recovered from the wound beds retained viable bacteria at an average density of 5 × 10⁸ CFU/mL, comparable to the initial encapsulation density (**Figure 6B, C**). Importantly, wound swabs collected from tissue beds directly beneath the recovered hydrogels yielded no detectable *A. faecalis* colonies across all animal replicates, supporting effective local biocontainment (**Figure 6B**). Furthermore, imaging of the recovered hydrogels revealed biofilm-like intra-gel accumulations resembling those observed *in vitro* (**Figure 6C**). Together, these findings demonstrated that hydrogel encapsulation supports prolonged bacterial persistence at the wound site while limiting detectable bacterial escape into the underlying wound bed, establishing a foundation for evaluating the therapeutic efficacy of these living materials in diabetic wound healing.

### Accelerated healing kinetics and histopathological analysis of healing skin in diabetic wounds treated with *A. faecalis*

Having established sustained bacterial persistence and local biocontainment within the wound environment, we next evaluated the therapeutic efficacy of *A. faecalis* cELM over time by monitoring wound closure in diabetic (*db/db)* mice. Wound closure was assessed throughout the 14-day treatment period and for an additional 7 days following treatment removal, allowing us to evaluate both the progression of healing during treatment and subsequent closure through day 21 (**Figure 7A, B**). During the initial healing phase (days 0–2), wound closure did not differ significantly amongst treatment groups. The first significant differences emerged on day 3, when wounds treated with freely suspended *A. faecalis* (AF Susp) exhibited greater closure than Blank Gel controls (*p* = 0.0449). By day 5, both *A. faecalis*-containing treatments (AF Susp and AF Gel) promoted significantly accelerated closure compared to Blank Gel control, with AF Susp also statistically outperforming PBS (**Figure 7A**). These improvements persisted through day 7, when both *A. faecalis*-containing treatments significantly enhanced closure relative to both vehicle controls.

**Figure 7.**
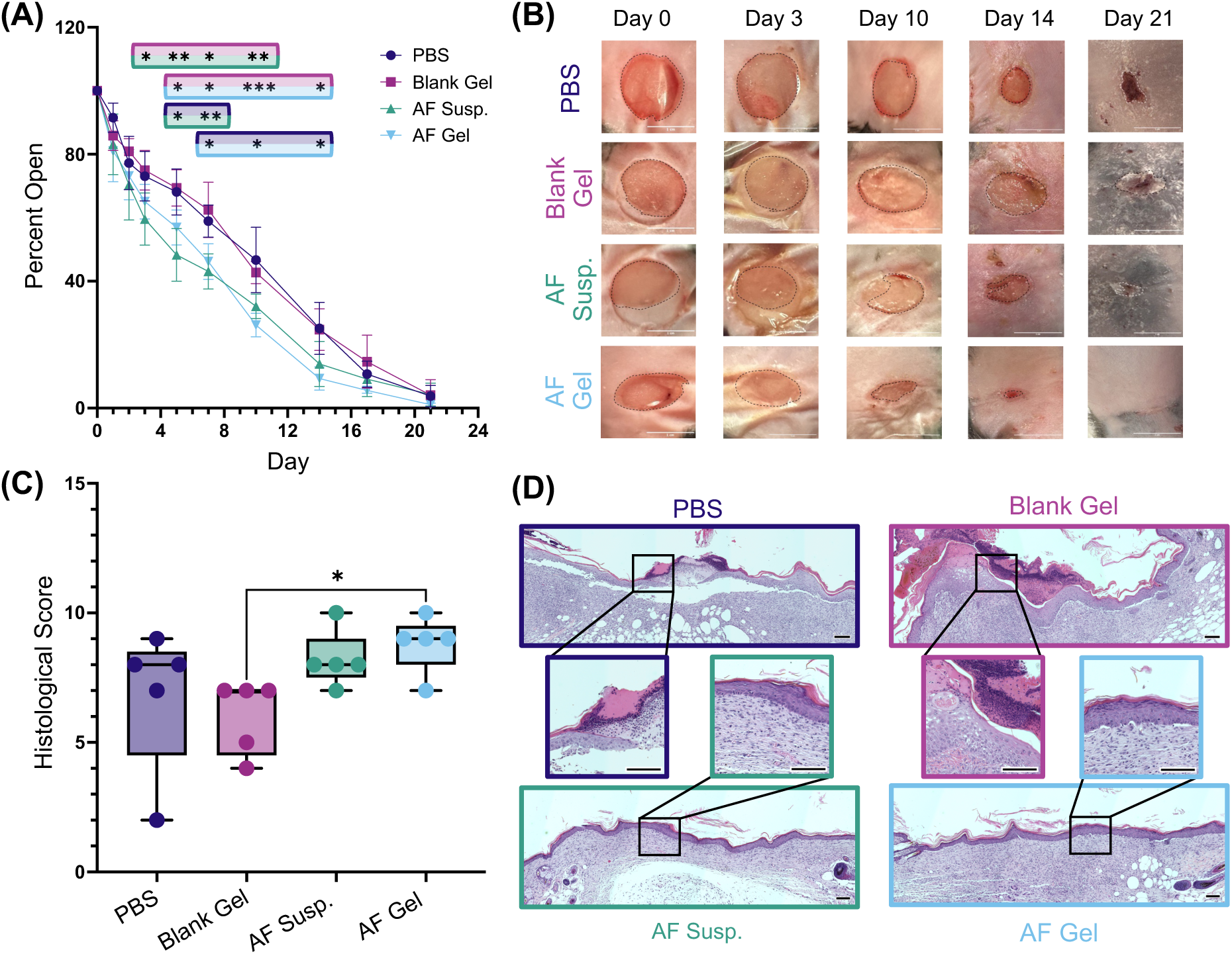
Macroscopic wound healing kinetics and quantitative histopathological regeneration. (A) Quantitative macroscopic wound closure tracking curves plotting the percentage of open wound area relative to day 0 over a 21-day healing timeline across four cohorts: sterile saline (PBS), blank vehicle hydrogels (Blank Gel), free *A. faecalis* suspensions (AF Susp), and active living hydrogels (AF Gel). Statistical markers highlighted with colors of groups being compared at each time point. (B) Images of wounds over 21-day experiment. Scale bar = 1 µm. (C) Histological SPOT score calculated for each condition based on closure, inflammation, and epithelial thickness. (D) Representative cross-sectional histopathological images of harvested wound beds sectioned through the wound center and stained with Hematoxylin and Eosin (H&E), illustrating advanced epidermal stratification and mature granulation tissue integration in the active AF Gel cohort compared to controls. For closure plot: data represent mean (N=5) ± standard deviation (SD) and *P ≤ 0.05; **P ≤ 0.01; ***P ≤ 0.001, ****P ≤ 0.0001 by 2-Way ANOVA with Tukey’s multiple comparison test. For histological score plot: data represent individual scores and *P ≤ 0.05 by Brown-Forsythe and Welch One-Way ANOVA with Dunnett’s T3 multiple comparisons test.

By day 10, treatment divergence occurred between the two active biotherapeutic delivery modes. Notably, the AF Gel maintained a significant healing advantage over both PBS and Blank Gel controls through day 14, whereas the AF Susp group no longer differed significantly from either control by day 14. Following treatment removal, wound closure converged across groups, with nearly complete macroscopic closure observed by day 21 as wounds were allowed to follow contractile healing common in mouse skin (**Figure 7A, B**). Overall, wounds treated with *A. faecalis* laden hydrogels showed significantly accelerated closure from day 5 through day 14, when treatments were removed, and unencapsulated suspension cultures lost their statistical healing advantage after day 10. Together, these findings demonstrated that both bacterial delivery approaches accelerated early wound closure, but hydrogel encapsulation extended the statistically detectable healing benefit through the full 14-day treatment period.

Although macroscopic wound closure converged across groups by day 21, differences in the quality of regenerated tissue may persist beyond apparent closure. To investigate this possibility, we conducted quantitative histopathological scoring of harvested day 21 wound tissues using the semi-quantitative SPOT grading matrix across five distinct parameters: *i)* Re-epithelialization, *ii)* Epidermal Thickness Index (ETI Score), *iii)* Keratinization, *iv)* Granulation Tissue (GT Score), and *v)* Scar Elevation Index (SEI Score) (**Figure 7C**) (*50*). The *A. faecalis*-laden hydrogel group exhibited the highest mean cumulative SPOT score (8.8 ± 1.1), followed by the free *A. faecalis* suspension (8.2 ± 1.1), PBS (6.8 ± 2.8), and Blank Gel (6.0 ± 1.4). These findings suggested a trend toward improved tissue restoration with bacterial treatment, particularly with hydrogel-mediated delivery.

Evaluation of individual histopathological parameters revealed additional differences in tissue architecture amongst treatment group (**Figure 7D, Table S3**). Notably, wounds treated with *A. faecalis*-laden hydrogels achieved the maximum re-epithelialization (REI = 2.0) and keratinization (KT = 2.0) scores, corresponding to 95–100% epidermal gap closure. Further, the AF Gel-treated wounds exhibited higher epidermal thickness (ETI = 1.8) and scar elevation (SEI = 1.8) scores relative to Blank Gel (ETI = 0.8, SEI = 0.6) and PBS controls (ETI = 1.0, SEI = 1.2) (**Table S2**). These elevated scores reflected restoration of normal epidermal thickness (95–105% of uninjured skin) and normal dermal contour heights in the AF Gel group compared to the epidermal hypoplasia (<95% thickness) and depressed scar contours observed in vehicle control groups (**Figure 7D, Table S2**). Granulation tissue scores confirmed that active treatments supported thick-to-intact dermal bed regeneration (GT = 1.2 for AF Gel and 1.6 for AF Susp), whereas Blank Gel controls retained thin, poorly organized granulation beds (GT = 0.8) (**Table S2**). Together, the macroscopic wound closure and histopathological findings demonstrated that an *A. faecalis* cELM promotes wound healing in a diabetic wound model, with sustained improvements in wound closure during treatment and histological features consistent with enhanced tissue restoration following treatment removal.

## DISCUSSION

Diabetic wounds present a critical challenge in healthcare characterized by persistent elevated proteolytic activity, leading to impaired keratinocyte migration (*11–22*). Although living cell therapies offer the capacity for continuous factor production in the wound, their application remains constrained by rapid clearance from the host and safety concerns surrounding uncontrolled microbial delivery (*3, 4, 23, 51*). Herein, we established a commensal Engineered Living Material (cELM) platform that achieves complete physical biocontainment of the commensal wound microbe *A. faecalis* while enabling continuous, localized release of its pro-healing secretome to accelerate wound healing. By systematically investigating the effect of polymer density and encapsulation density on metabolic activity and protein secretion over time, we established design rules for using synthetic PEG matrices encapsulating commensal *A. faecalis* as cELMs for secretome delivery and treatment of chronic wound environments (*1–4, 34, 35*).

To deploy living materials in wound care, a key requirement is biocontainment without preventing the transport of secreted therapeutic factors. We found that a clickable 4-arm 10 wt% PEG hydrogel with an estimated mesh size of about 5 nm allowed entrapment of *A. faecalis* cells while permitting cell function and viability, with the diffusion of nutrients into the gel and metabolic waste and therapeutic factors out of the gel (*36, 40, 52*). As demonstrated by total protein quantification and gel electrophoresis, the released secreted factors retained their molecular weight distribution when comparing suspension cultured cells and encapsulated cells. In contrast, higher polymer densities led to mesh sizes that hindered diffusion, whereas lower polymer densities had compromised physical biocontainment, potentially owing to defects network structure (*45, 46*). The sustained high viability of encapsulated *A. faecalis* within 10 wt% gels further supported the feasibility of this platform, while revealing the emergence of intra-gel biofilm-like accumulations within the hydrogel network.

Beyond physical mesh boundaries, the cellular microenvironment within the hydrogel matrix also plays an essential role in dictating long-term metabolic activity and secretory behavior. Counterintuitively, lower bacterial encapsulation densities (1 × 10^8^ CFU/mL) generated increased cumulative protein yields over extended culture compared to more densely seeded constructs (5 × 10^8^ and 9 × 10^8^ CFU/mL). This inverse density relationship illustrates that therapeutic secretome output is likely driven by per-cell metabolic efficiency. We hypothesize that spatial crowding and localized nutrient competition in high-density hydrogels stunt cellular kinetics over time. SDS-PAGE profiling confirmed that the 10 wt% PEG network readily permitted outward transport of the complete *A. faecalis* secretome without molecular weight cutoff limitations. Furthermore, bioactivity of the released secretome was confirmed to be similar in conditioned media from the low-density *A. faecalis* cELM and higher density suspension cultures in an *in vitro* scratch closure assay Crucially, the robust expansion of intra-gel aggregates and CV retention in lower-density formulations indicated that encapsulated bacteria actively produced biofilm-like secretions within the synthetic network (*1, 43, 44*).

Confocal imaging provided insight into the structural composition of intra-gel aggregates. Imaging revealed localized PI (red) signal adjacent to encapsulated cells, enclosed within a broader SYTO 9 (green) fluorescent envelope matching the physical boundaries of the matrix secretions seen in CV staining. The large, rounded objects in the stained gels likely reflect the binding properties of SYTO 9 and PI within biofilm architectures, as seen in literature (*53–56*). PI possesses a divalent positive charge (+2) leading to high electrostatic binding affinity, enabling it to selectively label dense, localized patches of extracellular DNA (eDNA) near cell surfaces, common in biofilms. SYTO 9, on the other hand, is an amphiphilic monovalent dye (+1) that exhibits broader matrix accessibility, binding across extracellular RNA, outer membrane vesicles, and some hydrophobic matrix proteins to illuminate the surrounding extracellular polymeric substance. This matrix deposition, coupled with the structural confinement of the PEG hydrogel, allows encapsulated commensal cells to function as stable, long-lasting bioreactors without escaping the carrier or compromising transport kinetics into the target tissue bed (*43*).

In translating this therapeutic system to an *in vivo* model, we saw biocontainment and viability of cells within the hydrogel matrix for 14 days in diabetic murine wounds, as well as the formation of biofilm-like intra-gel accumulations within the hydrogel matrix resembling those studied *in vitro* (*31*). Notably, the hydrogels shielded encapsulated cells from host clearance kinetics, maintaining complete physical confinement and high cell viability through day 14 without microbial escape into the wound bed. In contrast, topically applied unencapsulated bacteria experienced rapid host clearance, causing their localized presence and therapeutic impact to wane after day 10. This sustained retention drove accelerated tissue regeneration throughout the active application window. While unencapsulated suspension cultures lost their healing advantage as bacteria were cleared, *A. faecalis* cELMs maintained continuous, accelerated wound closure. The convergence of macroscopic wound closure across groups after day 14 reflects the transition to the late-stage, contraction-dominated healing characteristic of rodent skin models. Crucially, the therapeutic advantage established during the active treatment window resulted in superior histopathological SPOT scores at day 21, marked by complete re-epithelialization, restored normal epidermal thickness, and mature granulation tissue integration.

With the *A. faecalis* cELM platform and its relevance for wound healing demonstrated, avenues for future investigation include deep dives into mechanism for system optimization. For example, while bioactivity of the released secretome is evident, the specific active proteins, peptides, or metabolic factors responsible for enhancing healing remain unidentified (*29, 31*). The detailed proteomic and metabolic profiling could be used to elucidate the identity of such factors in future studies, with the potential to refine matrix design for promoting production and release of desired proteins within the secretome. Furthermore, probing the transcriptomic changes associated with the phenotypic shifts induced in *A. faecalis* by physical confinement and intra-gel biofilm formation could further guide platform refinement. Additionally, probing the dynamics of bacterial density, protein secretion, and tissue repair may further guide dosing and treatment time course. While murine models provide critical initial validation of biocontainment and bioactivity, rodent wounds heal predominantly through contraction rather than human-like re-epithelialization (*50, 57*); accordingly, future translation studies may move this platform forward into more complex model systems that better recapitulate human healing mechanics.

Together, these findings define key relationships between hydrogel network architecture, bacterial seeding density, and intra-matrix secretion dynamics for deploying commensal microbes as localized delivery depots. By ensuring complete physical biocontainment while sustaining cellular metabolic output and unhindered secretome diffusion, this platform overcomes the standard retention and safety barriers associated with live microbial therapies. This cELM framework offers a versatile bioengineering approach for leveraging host–commensal interactions to accelerate tissue regeneration in compromised wound environments.

## MATERIALS AND METHODS

### Hydrogel design

We prepared cELMs using bulk poly(ethylene glycol) (PEG) hydrogels via thiol-Michael addition crosslinking chemistry. We used four-arm PEG functionalized with vinyl sulfone (PEG-VS4, MW 5 kDa, Creative PEGWorks, PSB-4322) and four-arm PEG functionalized with thiol (PEG-SH4, MW 5 kDa, JenKem Technology USA, A7002-1). We maintained a stoichiometric balance of reactive end groups at a 1:1 ratio of thiol to vinyl sulfone groups (r = 1.0). We prepared precursor stock solutions of PEG-VS4 and PEG-SH4 in sterile 1× Dulbecco’s Phosphate-Buffered Saline (1× DPBS, pH 7.0, Gibco, 14-190-144). We calculated precursor volumes to yield a final polymer concentration of 10 wt% (and 5 wt% or 20 wt% where indicated) upon mixing and gelation. We immediately pipetted the polymer precursor solution into sterile rubber gaskets with an internal diameter of 8 mm and a thickness of 1 mm (total volume ∼50 µL) resting on sterile glass slides. Gelation proceeded at room temperature for ∼1.5 hours until a self-supporting, fully crosslinked network formed (*41*). We estimated the theoretical network mesh size (ξ) to be approximately 5 nm utilizing the Flory-Rehner relationships defined in Equation 1 and Equation 2 below (*36*).

### Bacterial strains and culture conditions

We incubated cultures of the commensal strain *Alcaligenes faecalis* (*A. faecalis*, strain maintained by Grice Lab at University of Pennsylvania) in tryptic soy broth (TSB, BD Difco, DF0370-17-3) at 37 °C on a shaker plate (150 rpm) overnight (*31*). We monitored cell growth through optical density measurements at 600 nm (OD600). To prepare cells for encapsulation, we harvested mid/late log-phase cultures via centrifugation at 4,000 rpm for 30 minutes at 4 °C. We washed and re-centrifuged the bacterial pellet with sterile 1× DPBS (pH 7.0) to remove residual growth media, repeating this process for a total of two washes. We determined cell density via serial dilutions and plating on tryptic soy agar with 5% sheep’s blood (Blood TSA) to correlate OD600 values with colony forming units per mL (CFU/mL).

### Encapsulation of bacteria within hydrogels

We gently mixed washed *A. faecalis* cell suspensions in 1× DPBS (pH 7.0) with the PEG precursor solutions to form cELM constructs. We adjusted the cell suspension density to achieve encapsulation densities of 1 × 10^8^, 5 × 10^8^, or 9 × 10^8^ CFU/mL within the final 10 wt% PEG hydrogels (5 wt% or 20 wt% hydrogels where indicated). We immediately cast the well-mixed precursor fluid into the 8-mm gaskets as described above. Once gelled, we carefully removed the bacteria-laden hydrogels from the gaskets and transferred them to sterile well plates containing nutrient media (M9 minimal media BD Difco, DF0485-17 supplemented with 0.5% tryptone Gibco, DF0123-17-3) for subsequent *in vitro* evaluations.

### Bacteria viability assays

We analyzed the survival and spatial distribution of encapsulated *A. faecalis* cells within the hydrogel network using the LIVE/DEAD BacLight bacterial viability kit (Invitrogen, L7012) after imaging on an Andor Dragonfly 600 confocal microscope system at 25x magnification. We incubated cell-laden hydrogels in 1× DPBS containing 3 µM SYTO 9 (green nucleic acid stain for all cells) and 15 µM propidium iodide (red nucleic acid stain for cells with compromised membranes) for 30 minutes. We stained non-bacteria-laden hydrogels under identical conditions to assess non-specific fluorophore binding to the polymer matrix.

We quantitatively tracked metabolic activity over a 14-day incubation period using PrestoBlue cell viability reagent (Invitrogen, A13261) across N = 4 independent hydrogel replicates per group. We placed hydrogels in fresh supplemented M9 minimal media in a well plate, added PrestoBlue reagent to a final concentration of 10% by volume, and incubated samples at 37 °C for 2 hours with shaking. Following incubation, we recorded the fluorescent signal corresponding to the reduction of resazurin to resorufin from the supernatant of the gels using a microplate reader (SpectraMax i3x, Molecular Devices, Delaware Biotechnology Institute).

### Polymer network model and transport through hydrogels

To establish the physical mechanisms governing secretome transport and bacterial containment, we applied Flory-Rehner network theory across varying polymer concentrations (5, 10, and 20 wt%) and bacterial encapsulation densities (0, 1 × 10⁸, 5 × 10⁸, and 9 × 10⁸ CFU/mL). Based on the equilibrium volumetric swelling ratio (*Q*) and the polymer volume fraction in the swollen state (*ν*₂,ₛ), we estimated the average hydrogel mesh size (ξ) via:

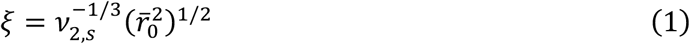

where (*r̄*₀²)¹ᐟ² represents the root-mean-square unperturbed end-to-end distance of the polymer chains between crosslinks, defined as:

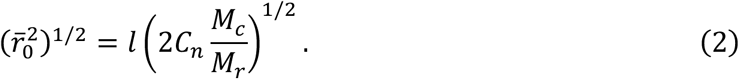

Here, *l* = 0.146 nm is the bond length along the PEG backbone, *C_n_* = 4.0 is the characteristic ratio for PEG, *M_r_* = 44 g/mol is the repeat unit molecular weight, and *M_c_* is the average molecular weight between crosslinks determined via Flory-Rehner equilibrium swelling analysis (*36, 41, 52*). In the absence of cells, increasing polymer concentration scaled the average network mesh from ξ ≈ 7.22 ± 0.13 nm for 5 wt% PEG gels to ξ ≈ 4.81 ± 0.07 nm for 10 wt% PEG gels, and ξ ≈ 2.69 ± 0.04 nm for 20 wt% hydrogels. When we encapsulated cells at up to 9 × 10⁸ CFU/mL, the estimated mesh sizes did not change significantly. This invariance demonstrated that bacterial loading did not compromise or alter matrix crosslinking (**Table S1**) (*36, 52*).

To evaluate how these networks discriminate between biological cargo and secreted factors of various sizes, we modeled relative diffusivity (*D_g_/D_0_*, where *D_g_* is the diffusion coefficient in the hydrogel and *D*₀ is the solute diffusivity in pure water). We estimated relative diffusivities using myoglobin (17 kDa, *r*ₛ ≈ 2.12 nm) and bovine serum albumin (BSA, 67 kDa, *r*ₛ ≈ 3.56 nm) as model secretome proteins, expressed as a function of swollen polymer volume fraction (*ν*₂,ₛ) and solute hydrodynamic radius (*r*ₛ):

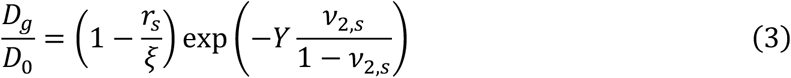

where *Y* represents the ratio of critical volume required for a translational solute jump to the free volume of water, taken as *Y* = 1 for water-swollen PEG networks (*36, 52*). Modeling the diffusion of BSA (67 kDa) in 20 wt% gels showed that the mesh size was smaller than the hydrodynamic radius of the protein, leading to the total hindrance of large protein diffusion through the matrix. Conversely, the 5 wt% formulation permitted robust diffusion due to a mesh size significantly larger than BSA; however, its looser network and inherent structural heterogeneities heightened concerns regarding long-term biocontainment, bacterial escape, and uncontrolled protein delivery. The 10 wt% gels (ξ ≈ 4.81 nm) resolved these concerns by maximizing biocontainment while maintaining an optimal transport window, permitting high diffusivity ratios of *D_g_/D_0_* ≈ 0.504 for myoglobin (17 kDa, *D_g_* ≈ 5.7 × 10⁻¹⁰ m²/s) and *D_g_/D_0_* ≈ 0.234 for BSA (*D_g_* ≈ 2.09 × 10⁻¹¹ m²/s). Because single *A. faecalis* cells (*r*ₛ ≈ 500 nm) exceed the 10 wt% mesh dimensions by two orders of magnitude (*r*ₛ ≫ ξ), their theoretical diffusivity drops to *D_g_* → 0. This provides a physical mechanism for complete bacterial confinement paired with unhindered secretome diffusion (**Table S3**) (*36, 52*).

### Protein release assay and quantification

To characterize secretome release kinetics through the hydrogel network, we incubated bacteria-laden hydrogels (N = 4 independent hydrogel replicates per group) in sterile supplemented M9 minimal media (pH 7.0) at 37 °C. We collected supernatants daily and replaced them in full with fresh media to maintain sink conditions and ample nutrients. To eliminate potential debris, we centrifuged collected supernatants at 4,000 rpm for 30 minutes at 4 °C and passed them through a sterile 0.22 µm polyether sulfone (PES) filter. We quantified total protein content within sterile-filtered supernatants using the Pierce Rapid Gold BCA protein assay kit (Fisher Scientific, A55862) according to the manufacturer’s microplate protocol.

To evaluate the composition and molecular weight distribution of the released secretome, we performed sodium dodecyl sulfate-polyacrylamide gel electrophoresis (SDS-PAGE) on hydrogel supernatants using precast polyacrylamide gels (Bio-Rad Laboratories, Inc., 456-1096). We visualized protein bands using SYPRO Ruby protein gel stain (Invitrogen, S11791) and compared them directly against supernatants from liquid culture suspensions of *A. faecalis* at an equivalent concentration.

### Conditioned media preparation and characterization

We prepared conditioned media to evaluate the biological activity of the *A. faecalis* secretome from cells grown in suspension and those encapsulated. We incubated bacteria-laden hydrogels (1 × 10^8^, 5 × 10^8^, or 9 × 10^8^ CFU/mL) in sterile supplemented M9 minimal media for 24 hours at 37 °C to allow secretome accumulation. We collected the resulting supernatant, centrifuged it at 4,000 rpm for 30 minutes at 4 °C, and sterile-filtered it through a 0.22 µm filter to ensure complete removal of bacterial cells or debris. We prepared control conditioned media under identical incubation and filtration conditions using blank 10 wt% PEG hydrogels (vehicle control) and unencapsulated *A. faecalis* suspension cultures (positive control). To evaluate biological effects on mammalian cells, we mixed these sterile conditioned media stocks at a 10% (v/v) concentration into standard keratinocyte culture media (Lonza, CC-3111) for downstream functional assays (*31*).

### Biofilm detection staining and imaging

We characterized intra-gel biofilm development of *A. faecalis* entirely *in vitro* utilizing a dedicated set of hydrogels (N = 4 independent hydrogel replicates per condition). We cultured newly fabricated gels *in vitro* and harvested them at designated experimental timepoints. We fixed gels in ice-cold ethanol and subsequently stained them with 0.1% crystal violet (CV) solution. We performed qualitative biofilm imaging using an EVOS M7000 Imaging System (ThermoFisher Scientific). To quantitatively evaluate biofilm formation, we resolubilized bound crystal violet by incubating stained gels in 70% ethanol for two hours at room temperature on a shaker plate at 150 rpm. We subsequently measured the absorbance of the resolubilized dye using a microplate reader (*43, 53–56*).

### Scratch closure assay

We seeded primary human epithelial keratinocytes onto tissue culture plastic and cultured them to form a confluent monolayer. We created a cell-free scratch wound in the confluent monolayer of each well using a sterile 200 µL pipette tip. Following scratch generation, we washed wells with sterile 1× DPBS to remove cellular debris and dislodged cells. We treated the scratched keratinocyte monolayers with standard keratinocyte culture media supplemented with 10% (v/v) of the respective sterile filtered conditioned media group: *A. faecalis*-laden bulk hydrogel conditioned media (from encapsulation density: 1 × 10^8^ CFU/mL or 5 × 10^8^ CFU/mL), *A. faecalis* suspension culture conditioned media (collected after 9 hours of culture and 18 hours of culture), or vehicle control (PBS or blank hydrogel) conditioned media. Based on results obtained from total protein quantification, the total protein after each 24-hour fraction was estimated to be about the same concentration. Thus, the fractions from each 24-hour period were combined to make a solution containing the secreted factors from all 14 days of collection.

We imaged the initial scratch area immediately after treatment application (t = 0 hours) using light microscopy, recording spatial coordinates within each well. We incubated plates at 37 °C in a humidified incubator containing 5% CO2. After a 24-hour incubation period, we re-imaged scratch wounds at identical spatial coordinates (t = 24 hours). We quantified wound gap closure from digital images utilizing the automated ImageJ plugin “Wound healing size tool, updated” and reported it as percent closure relative to t = 0 (*49*).

### Animal models and husbandry conditions

All animal experiments were conducted in compliance with protocols approved by the Institutional Animal Care and Use Committee (IACUC) at the University of Delaware (UD, Protocol AUP #1229). We utilized male Lepr^db (*db/db*) mice (8 weeks of age), a genetically diabetic mouse model of type 2 diabetes and impaired wound healing (The Jackson Laboratory, B6.BKS(D)-Leprdb/J, 000697). We housed mice individually in a temperature- and humidity-controlled facility at the University of Delaware on a 12-hour light/dark cycle with access to food and water as needed (*31, 57*).

### Excisional wounding and treatment of murine diabetic wounds

Prior to surgical procedures, we anesthetized mice via isoflurane inhalation. We shaved the dorsal region and depilated the area using Nair hair remover lotion. We created two full-thickness excisional skin wounds (8 mm in diameter) on the shaved dorsum on opposite sides of the midline using a standard punch biopsy tool (Miltex, 8 mm). We positioned wounds at a fixed distance to prevent physical or therapeutic cross-contamination. We randomly assigned and treated wounds across four experimental cohorts (N = 5 mice per group): *A. faecalis*-laden bulk PEG hydrogel discs (5 × 10^8^ CFU/mL final encapsulation density), a free *A. faecalis* suspension bolus (same volume as hydrogel disc at 5 × 10^8^ CFU/mL in sterile saline applied topically), blank PEG hydrogel discs as a vehicle control, or sterile saline as an untreated vehicle control. Following treatment application, we covered wounds with an optically clear, sterile dressing (3M™ Tegaderm™ Transparent Film Roll, 7100223172) to secure treatments and maintain moisture. We tracked wound closure through daily photographs on days 0, 1, 2, 3, 5, 7, 10, 14, 17, and 21 taken using an IVIS Lumina III *in vivo* Imaging System (Delaware Center for Musculoskeletal Research). We quantified wound area from digital images using ImageJ while blinded to treatment groups (*31, 57*).

### Bacterial CFU quantification of wounds

To evaluate the *in vivo* persistence, viability, and biocontainment of delivered bacterial gels and boluses (N = 5 mice per group), we thoroughly swabbed wound beds on day 14. We plated swabs collected from all hydrogel-treated wounds on nutrient agar to screen for escaped cells or native microbial colonization; we observed no observable *A. faecalis* growth across these plates, confirming successful physical containment within the PEG network. We similarly plated swabs collected from the free suspension group, which yielded minimal to no observable growth, confirming rapid clearance of unencapsulated bacteria from the wound bed (*31, 57*)

### Histological sectioning and analysis

Following the designated *in vivo* experimental endpoint (day 21), we manually harvested wound tissue beds along with surrounding normal skin margins (N = 5 mice per group). We immediately fixed collected tissue samples in 4% paraformaldehyde (PFA) in PBS at 4 °C to preserve structural integrity. Fixed tissue specimens were transferred to the Histology Core at Nemours Children’s Health for processing, embedding, sectioning, and routine staining. Skin samples were processed using standard automated protocols, embedded in paraffin blocks, and sectioned through the center of the wound bed to capture cross-sections at maximum diameter. They were mounted sections onto glass slides and stained with Hematoxylin and Eosin (H&E) to visualize overall tissue architecture, cellular infiltration, and epidermal morphology.

We performed quantitative histopathological evaluation of wound healing blinded using the semi-quantitative SPOT grading matrix adapted from van de Vyver *et. al.* We captured representative images of wound tissue sections under brightfield microscopy using an EVOS M7000 Imaging System. Using ImageJ, we graded tissue regeneration individually across five structural indices: Re-epithelialization, Epidermal Thickness Index (ETI Score), Keratinization, Granulation Tissue (GT Score), and Scar Elevation Index (SEI Score), summing them to determine the final Total SPOT Score (*50*).

### Microscopy and image processing

We performed fluorescence and confocal imaging of stained hydrogels using an Andor Dragonfly 600 confocal microscope system. For 3D bacterial viability imaging, we collected 50-µm tall z-stacks using the LIVE/DEAD BacLight stain. To minimize bias, we established and maintained standard laser power and gain settings across all experimental groups. We performed quantitative bacterial cell counting from confocal z-stack images via 3D spatial coordinate modeling utilizing the “Spots” object detection tool in Imaris software. We imaged keratinocyte cell monolayers using an EVOS M7000 Imaging System (*56*).

### Statistical analysis

We executed all statistical analyses using GraphPad Prism (version 11). All quantitative data are expressed as mean ± standard deviation (SD). *P*-values < 0.05 were considered statistically significant (*P* < 0.05, \**P* < 0.01, *P* < 0.001, \*\*\**P* < 0.0001). Specific sample sizes (N), *P*-values, and statistical tests (including post-hoc multiple comparison tests) performed for each experiment are explicitly detailed within the respective figure legends and main text. Sample sizes for *in vitro* assays and *in vivo* animal cohorts were selected based on preliminary studies to provide sufficient statistical power (β = 0.80, α = 0.05).

## Acknowledgments

The authors would like to thank Christina Stinger for her assistance with the surgical procedures for the murine excisional wound healing studies. We express our gratitude to Ellen K. White, MD, PhD, Simon Knight, PhD, and Jamie Ting-Chun Pan, PhD, from the Grice Lab at the University of Pennsylvania for training on the scratch assay protocol, and Luisa Palmese, PhD, from the University of Delaware for training on hydrogel synthesis. Additionally, the authors thank Jeffrey Caplan, PhD, at the Delaware Biotechnology Institute Bioimaging Center for training on the Andor Dragonfly. The authors acknowledge the use of BioRender.com (2026) in all figures.

## Author contributions

A.M.K., M.O.S., E.A.G., and C.O.L. conceptualized the study. C.O.L. developed the methodology, conducted the investigation, performed data visualization, and wrote the original manuscript draft. A.M.K. and M.O.S. supervised the project. All authors (C.O.L., M.O.S., A.M.K., and E.A.G.) participated in reviewing and editing the manuscript

## Data, code, and materials availability

All data needed to evaluate and reproduce the results in the paper are present in the paper and/or the Supplementary Materials. This study did not generate new materials or unique code. Biological materials and reagents are commercially available or can be obtained from the corresponding author upon reasonable request.

