## Supplemental Material for "Commensal Engineered Living Materials for Sustained Secretome Delivery and Wound Healing"

Claire O. Lois *et al.*

**This PDF file includes:**

Figs. S1 to S5
Tables S1 to S3

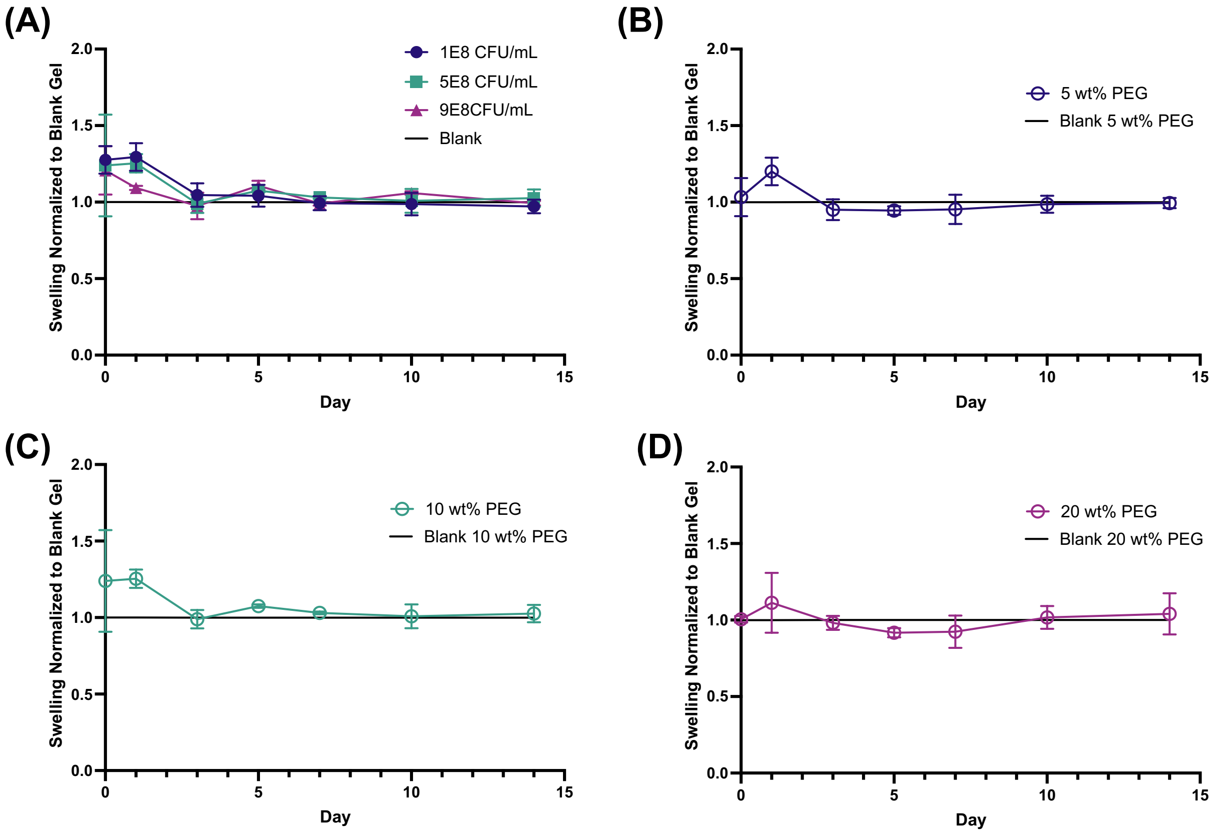


**Fig. S1. Normalized hydrogel swelling over time as a function of bacterial encapsulation density and polymer concentration.** (A) Normalized swelling ratio of 10 wt% PEG hydrogels encapsulated with varying bacterial densities ($1 \times 10^8$, $5 \times 10^8$, and $9 \times 10^8$ CFU/mL) over 14 days. (B–D) Normalized swelling ratio of hydrogels with a constant bacterial encapsulation density of $5 \times 10^8$ CFU/mL across different polymer weight percentages: (B) 5 wt% PEG, (C) 10 wt% PEG, and (D) 20 wt% PEG. In all panels, swelling volume is normalized to the corresponding unladen (blank) hydrogel of the same polymer weight percentage, represented by the solid black horizontal line ($y = 1.0$). Data are presented as mean $\pm$ SD.

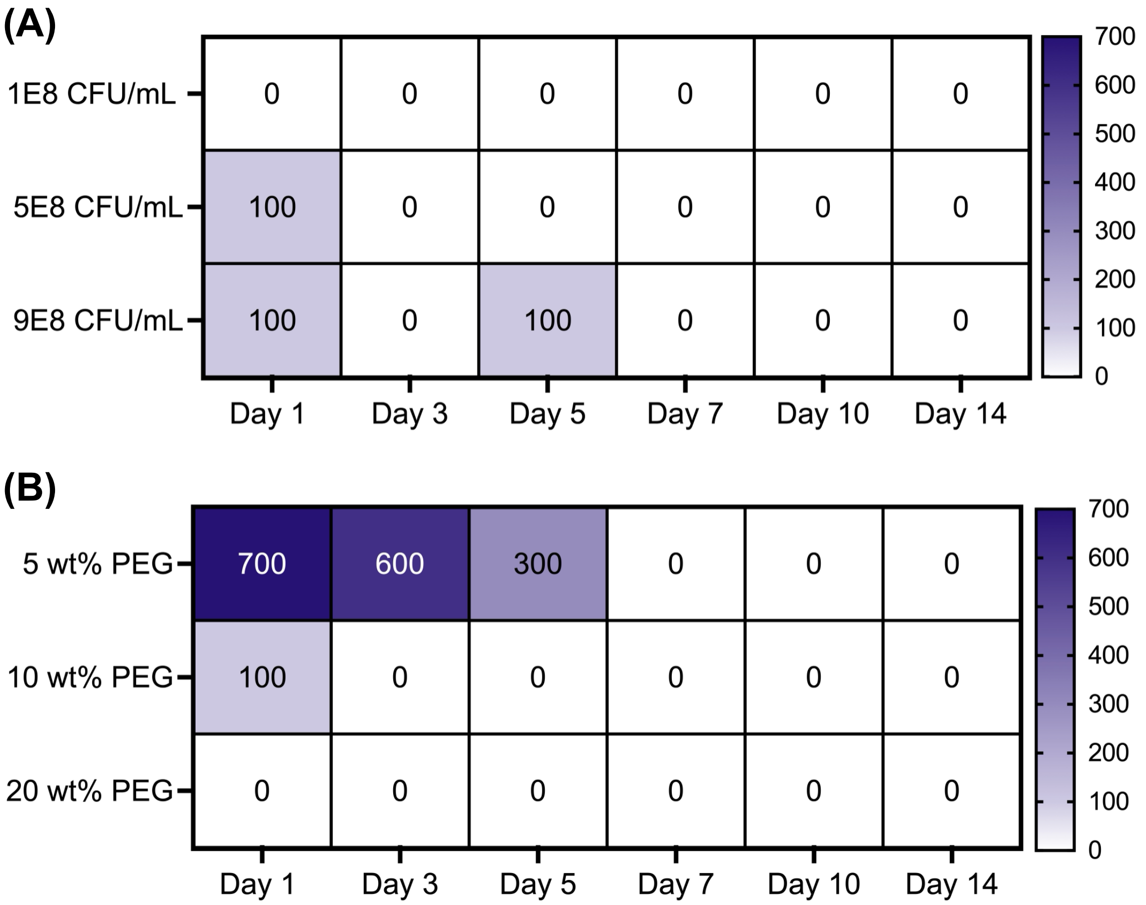


**Fig. S2. Evaluation of cell escape from hydrogels *in vitro*.** Heatmaps quantifying bacterial escape into supernatant over 14 days. (A) Escape kinetics across varying bacterial encapsulation densities in 10 wt% hydrogels. (B) Escape kinetics across varying PEG polymer concentrations (5-20 wt%) at fixed bacterial encapsulation. Color scale corresponds to escaped CFU count per mL.

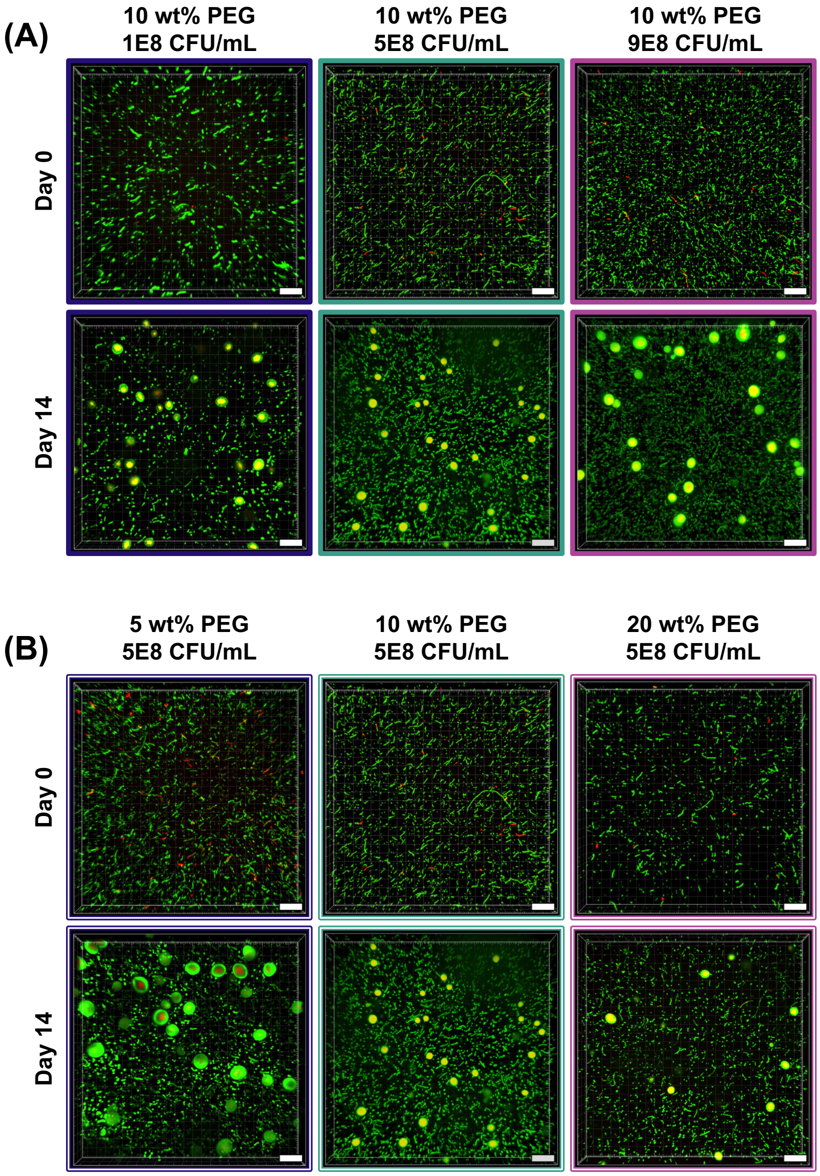


**Fig. S3. 3D images of hydrogel constructs at Day 0 and Day 14.** Full 3D spatial distribution and gel integrity of *A. faecalis* cELMs over 14 days. Z-stack 3D volume renderings depicting whole hydrogel constructs at initial encapsulation (Day 0) and following 14 days of culture (Day 14). Scale bars are 70 µm. (A) Variation of encapsulation density for *A. faecalis* and (B) variation of hydrogel density.

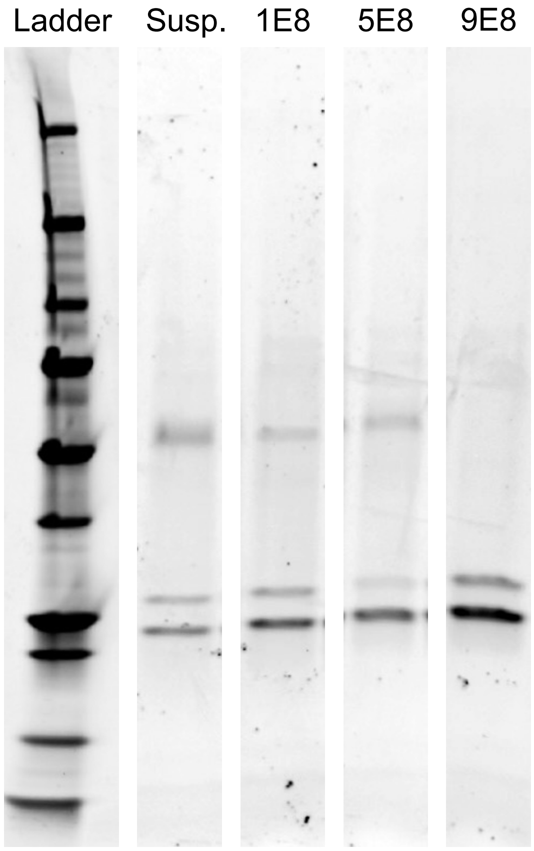


**Fig. S4. SDS-PAGE profile and densitometric analysis of conditioned media.** Representative SDS-PAGE gel images. Lanes display a molecular weight marker (Ladder), conditioned media from an 18 h cell-free suspension (Susp.), and conditioned media pooled over 14 days from hydrogel constructs encapsulated at cell densities of $\mathbf{1} \times \mathbf{10^8}$ CFU/mL (1E8), $\mathbf{5} \times \mathbf{10^8}$ CFU/mL (5E8), and $\mathbf{9} \times \mathbf{10^8}$ CFU/mL (9E8).

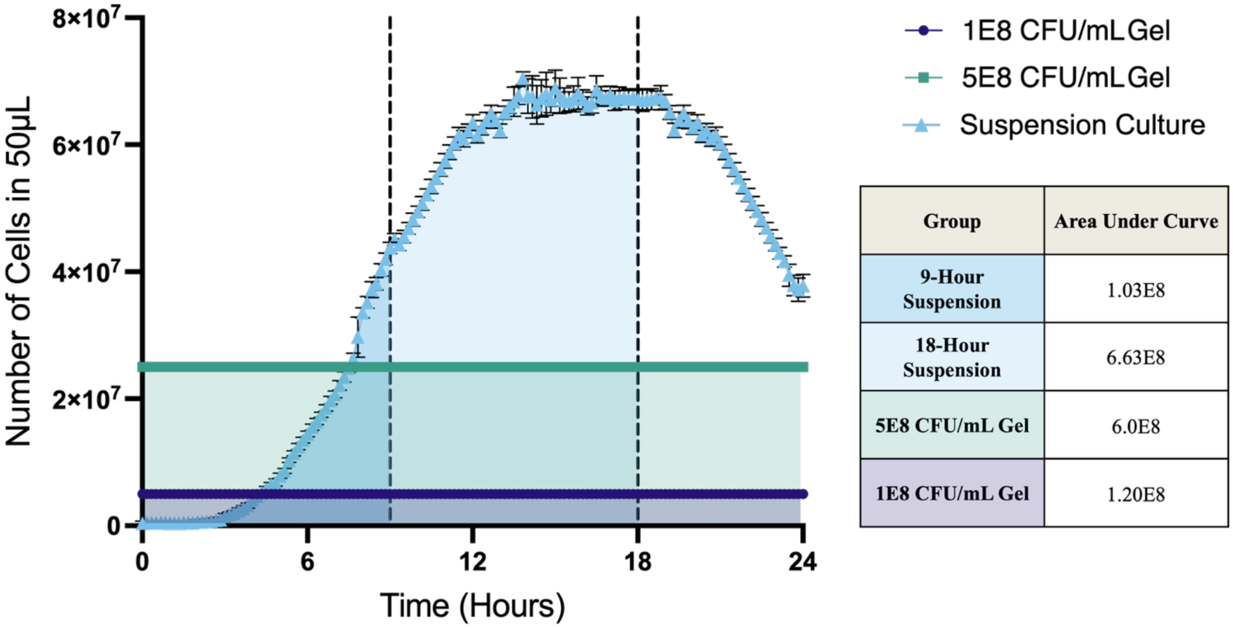


| Group | Area Under Curve |
|---|---|
| 9-Hour Suspension | 1.03E8 |
| 18-Hour Suspension | 6.63E8 |
| 5E8 CFU/mL Gel | 6.0E8 |
| 1E8 CFU/mL Gel | 1.20E8 |

**Fig. S5. Time-integrated cell number analysis for selecting suspension culture controls matched to hydrogel conditions.** Comparison of cell accumulation over time in growing *A. faecalis* suspension culture versus constant-cell hydrogel conditions within a representative $\mathbf{50}\ \boldsymbol{\mu}$L volume. Suspension culture cell numbers were integrated over time (shaded regions) to serve as a quantitative proxy for relative secretome production capacity. Hydrogel cell populations ($\mathbf{1 \times 10^8}$ and $\mathbf{5 \times 10^8}$ CFU/mL) were assumed constant over 24 h based on experimental measurements. The area under the curve (AUC) values (right table) show that integrating suspension cell growth over 0–9 h (AUC = $\mathbf{1.03 \times 10^8}$) and 0–18 h (AUC = $\mathbf{6.63 \times 10^8}$) yields time-integrated cell numbers comparable to the 24 h constant $\mathbf{1 \times 10^8}$ CFU/mL (AUC = $\mathbf{1.20 \times 10^8}$) and $\mathbf{5 \times 10^8}$ CFU/mL (AUC = $\mathbf{6.0 \times 10^8}$) hydrogel conditions, respectively. Consequently, 9 h and 18 h suspension time points were selected as matched controls for subsequent secretome analyses.

**Table S1. Estimated mesh sizes.**

| Group | 5 wt% PEG | 10 wt% PEG | 20 wt% PEG |
|---|---|---|---|
| **0 CFU/mL** | 7.22 $\pm$ 0.13 nm | 4.81 $\pm$ 0.07 nm | 2.69 $\pm$ 0.04 nm |
| **1E8 CFU/mL** | | 4.90 $\pm$ 0.15 nm | |
| **5E8 CFU/mL** | 7.09 $\pm$ 0.18 nm | 4.79 $\pm$ 0.13 nm | 2.64 $\pm$ 0.11 nm |
| **9E8 CFU/mL** | | 4.75 $\pm$ 0.19 nm | |

**Estimated hydrogel mesh sizes as a function of polymer concentration and bacterial encapsulation density.** Average network mesh sizes ($\xi$, nm) estimated using Flory-Rehner equilibrium swelling theory across varying PEG hydrogel weight percentages (5, 10, and 20 wt%) and bacterial encapsulation densities ($0$, $1 \times 10^8$, $5 \times 10^8$, and $9 \times 10^8$ CFU/mL). Mesh sizes scaled inversely with polymer concentration while remaining invariant across increasing cell densities, confirming that bacterial encapsulation does not disrupt network crosslinking. Data are presented as mean $\pm$ standard deviation.

**Table S2. SPOT histological scoring**

| Group | ID | REI | ETI | KT | GT | SEI | Total |
|---|---|---|---|---|---|---|---|
| AF | 1R | 2 | 2 | 2 | 2 | 2 | 10 |
| | 2R | 2 | 1 | 2 | 1 | 1 | 7 |
| | 3L | 2 | 1 | 2 | 1 | 2 | 8 |
| | 7L | 2 | 1 | 2 | 2 | 1 | 8 |
| | 8R | 2 | 1 | 2 | 2 | 1 | 8 |
| | **Average** | 2 | 1.2 | 2 | 1.6 | 1.4 | 8.2 |
| AF Gel | 2L | 2 | 2 | 2 | 1 | 2 | 9 |
| | 4L | 2 | 2 | 2 | 1 | 2 | 9 |
| | 6L | 2 | 2 | 2 | 2 | 2 | 10 |
| | 7R | 2 | 1 | 2 | 1 | 1 | 7 |
| | 9R | 2 | 2 | 2 | 1 | 2 | 9 |
| | **Average** | 2 | 1.8 | 2 | 1.2 | 1.8 | 8.8 |
| PBS | 1L | 2 | 1 | 2 | 2 | 2 | 9 |
| | 4R | 2 | 1 | 2 | 2 | 1 | 8 |
| | 5R | 2 | 1 | 2 | 1 | 1 | 7 |
| | 8L | 1 | 1 | 0 | 0 | 0 | 2 |
| | 10L | 2 | 1 | 2 | 1 | 2 | 8 |
| | **Average** | 1.8 | 1 | 1.6 | 1.2 | 1.2 | 6.8 |
| Blank Gel | 3R | 2 | 0 | 2 | 2 | 1 | 7 |
| | 5L | 2 | 1 | 2 | 1 | 1 | 7 |
| | 6R | 1 | 1 | 2 | 0 | 0 | 4 |
| | 9L | 2 | 1 | 2 | 0 | 0 | 5 |
| | 10R | 2 | 1 | 2 | 1 | 1 | 7 |
| | **Average** | 1.8 | 0.8 | 2 | 0.8 | 0.6 | 6 |

**Semi-quantitative SPOT histological scoring of wound healing at day 21.** Blinded histopathological evaluation of harvested wound tissue sections across treatment groups ($N = 5$ mice per group) using the semi-quantitative SPOT grading matrix. Individual animal scores (ID) and group averages are reported for five structural indices: re-epithelialization index (REI), epidermal thickness index (ETI), keratinization (KT), granulation tissue maturity (GT), and scar elevation index (SEI). Individual category scores were summed to yield the final Total SPOT Score for each specimen, evaluating overall tissue regeneration, epidermal reconstruction, and wound resolution relative to controls.

**Table S3. Estimated solute diffusivity.**

| Group | Model Solute | $D_g/D_0$ | $D_g$ |
|---|---|---|---|
| **5 wt% PEG** | Myoglobin, 17 kDa | 0.683 | 1.02E-10 $m^2/s$ |
| | BSA, 67 kDa | 0.490 | 4.36E-11 $m^2/s$ |
| **10 wt% PEG** | Myoglobin, 17 kDa | 0.504 | 7.57E-11 $m^2/s$ |
| | BSA, 67 kDa | 0.234 | 2.09E-11 $m^2/s$ |
| **20 wt% PEG** | Myoglobin, 17 kDa | 0.156 | 4.36E-11 $m^2/s$ |
| | BSA, 67 kDa | -- | 2.33E-11 $m^2/s$ |

**Estimated relative diffusivities ($D_g/D_0$) and diffusion coefficients ($D_g$) for model secretome proteins across hydrogel formulations.** Modeled transport parameters for myoglobin (17 kDa, $r_s \approx 2.12$ nm) and bovine serum albumin (BSA, 67 kDa, $r_s \approx 3.56$ nm) within 5, 10, and 20 wt% PEG hydrogel networks. $D_g/D_0$ represents the solute diffusivity within the gel relative to pure water, and $D_g$ represents the absolute diffusion coefficient in $m^2/s$.
